# DEEPNET-LLM: Highly accurate structure prediction of unknown metabolites from mass spectra via their molecular neighborhoods

**DOI:** 10.64898/2026.10.28.762558

**Authors:** Alexey V Melnik, Alexander Semenov, Andrew Roberts, Dana Moradi, Elizabeth A Coler, Jennifer Goldstein, Bibhusha Ojha, Nathan J Swift, Suada Demirovic, Mark Peczuh, Thomas D Horvath, Melinda A Engevik, Vladimir Boginski, Alexander A Aksenov

## Abstract

Most molecular features detected by untargeted mass spectrometry-based metabolomics remain unannotated. This so-called “dark matter” currently comprises more than 90% and likely holds uncharacterized bioactive chemistry. Structure prediction from fragmentation spectra remains an outstanding challenge, because a spectrum does not uniquely determine a structure and the space of candidate structures is mostly unbounded; correspondingly, existing methods rarely return the correct *de novo* structure as their top prediction. Here we describe *De novo* Ego network-Enabled Propagation and Neighborhood Extrapolation of Topology using Large Language Models (DEEPNET-LLM) reasoning approach, which uses Molecular Community Networking (MCN) to restrict the search of a query spectrum to its local neighborhood within the global metabolome, and applies AI reasoning to propagate structural information from neighbors to the query. For a collection of 900 authentic standards (2,249 spectra) that have never been publicly released to exclude possibility of data leakage, DEEPNET-LLM returned the correct structure as its top prediction for 84.7% of spectra. Accuracy was independent of precursor mass across a range over which the number of candidate structures grows by 30 orders of magnitude, and was comparable across ion polarity, adducts and in-source fragmentation. On a random subset of the public MassSpecGym benchmark, DEEPNET-LLM reached 85.9% top-1 accuracy excluding stereochemistry. We showcase the use of DEEPNET-LLM with an example of a discovery of tryptamine amidate of a bile acid in mouse gut, confirmed against a synthetic standard. The DEEPNET-LLM approach is orthogonal to existing *in silico* methods and can be combined with them.

**One-sentence summary:** Restricting the search to the molecular neighborhoods of the global metabolome enables accurate structure prediction for unknown metabolites from their mass spectra.

## Introduction

Biological processes are largely regulated by small molecules. Metabolites, signaling lipids, hormones and microbial natural products can bind to protein targets as agonists or antagonists, and thereby regulate enzyme activity, gate ion channels and initiate signaling cascades. Understanding and manipulating biology, including the design of therapeutics, therefore requires knowledge of both the protein target and the small molecules that act upon it. Molecules made by living systems are of particular importance: because evolution has selected them for compatibility with cellular chemistry, natural products and their derivatives remain the dominant source of therapeutics(*1*). The structural diversity of biological chemistry is large, and its extent is currently not known.

The protein part of the problem has now been largely resolved. Protein structure prediction was intractable for decades, because the conformational space available to a polypeptide chain is far too large to enumerate, and structures were determined experimentally, one at a time. AlphaFold resolved the problem by learning the regularities of folded structure from known examples, which constrained the space to be searched rather than exhausting it(*2*). The complementary problem, identifying the structure of small molecules, remains open.

Mass spectrometry is the most capable tool available for exploring the chemistry of living systems. It detects analytes at attomole to femtomole levels across a wide dynamic range (typically 3-5 order of magnitude for intrascan dynamic range for most commercial instruments), requires no prior knowledge of what is present and, through tandem fragmentation (MS/MS, and higher order fragmentation, MS^n^), can be used to determine structure. Coupled to liquid or gas chromatography, or even capillary electrophoresis, MS resolves tens of thousands of molecular features in a single untargeted acquisition(*3*). Most of these features, however, cannot be assigned a structure. This comprises so-called “dark matter” of metabolomics(*4*), and it has been reduced over the past decade, with annotation rates rising from under 1% to approximately 8%. This was achieved through the growth of reference libraries, discovery by molecular networking and repository-scale search(*5–7*), but the large majority of detected features remain unannotated.

Two routes to closing this gap have been pursued, collecting reference data and *in silico* predictions. Reference libraries require authentic standards, so they grow only as fast as compounds can be synthesized and measured. They are also biased toward commercially available synthetic compounds, many of which do not occur in living systems, so library growth does not translate proportionally into biological coverage. *In silico* structure prediction has been advancing rapidly. Learned spectral representations have sharpened the similarity measure underlying networking (MSBERT; DreaMS)(*8*, *9*), fingerprint prediction has refined structural inference from fragmentation (MIST)(*10*), and consensus scoring has propagated *in silico* annotations to networked neighbors (ConCISE)(*11*). Nevertheless, on the MassSpecGym benchmark none of the baseline *de novo* generation methods returned a correct structure at either top-1 or top-10 (*12*), and a recent leakage-controlled reassessment places the best subsequent *de novo* methods at 4.1% top-10 accuracy, with most top-1 accuracies between 0% and 2%(*13*). Top-10 lists are of limited use to biologists, who need a single correct structure to interpret an experiment. Retrieval against structural databases performs substantially better, but cannot propose structures that are not already catalogued. SIRIUS with CSI:FingerID, the most widely used tool of this class, ranks candidates drawn from existing databases(*14*, *15*). Retrieval-guided generation, in which MetGenX conditions a transformer decoder on spectrally retrieved reference structures, improves candidate retrieval substantially, but its gain in *de novo* prediction on structure-disjoint benchmarks remains modest (*16*).

Small molecule structure prediction from mass spectra is thus in a position similar to protein structure prediction before AlphaFold: the task is underdetermined because it is posed against a space that is unbounded and of unknown size. Here, we propose to constrain the candidate set before prediction is attempted, rather than searching chemical space at large. To achieve this, we leverage Molecular Community Networking (MCN), which enables placing each spectrum into a molecular context among its closest counterparts(*17*). MCN applies a molecular family-specific connectivity strategy that organizes the network into contiguous molecular communities, and it has been used to build a global network across all public GNPS data(*17*). MCN thus overcomes the key limitation of the conventional molecular networking which applies a single, uniform similarity threshold(*3*, *5*); because fragmentation behavior varies widely across chemical space, this fractures the connectivity of many molecular families and does not correctly describe molecular connectivity patterns. MCN-based networking makes it possible to place an arbitrary query spectrum within its global molecular neighborhood. We show that agentic AI-assisted reasoning over this neighborhood allows predicting complete structures for most query spectra, and that its accuracy does not degrade as the space of candidate structures grows. We termed this approach *De novo* Ego network-Enabled Propagation and Neighborhood Extrapolation of Topology using Large Language Models (DEEPNET-LLM).

## Results

### Constructing the propagation neighborhood of a query spectrum

DEEPNET-LLM begins by placing the query spectrum among the spectra it most resembles. The 8.4 million clustered spectra of public GNPS data are indexed with Hierarchical Navigable Small World (HNSW)-MS, an approach that utilizes pre-constructed hierarchical graph to retrieve nearest neighbors in logarithmic rather than linear time(*18*). The *k* most similar spectra (*k* = 25 throughout; the parameter is adjustable) constitute the ego network of the query (Figure 1).

**Figure 1.**
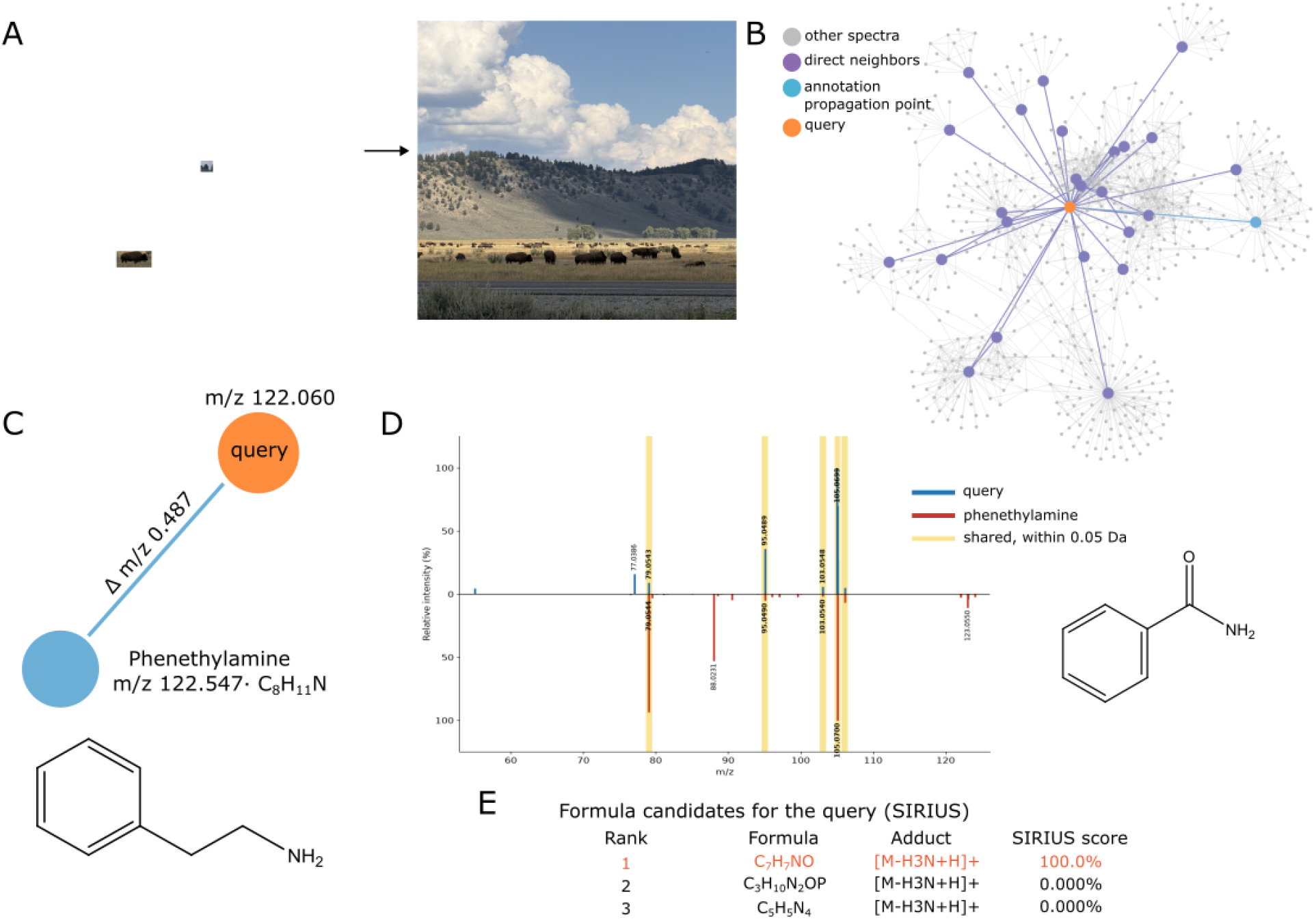
A conceptual explanation of the DEEPNET-LLM approach. **a)** Importance of context in annotation. (Left) In isolation, two features are very difficult to identify unambiguously. In context of the whole picture (Right), the same features can be confidently identified as a buffalo and a tree, correspondingly. b) An example of a propagation neighborhood used for annotation propagation, showing the query spectrum, the ego network of its k = 25 first-order spectral matches, and the outward expansion to second- and higher-order neighbors recruited from the MCN, drawn here to two hops. One node and its edge is highlighted in blue to show how the information for it is propagated, as an example. c) Annotation propagation for the node and edge highlighted in blue on panel (b) via comparison to the annotated neighbors from which propagation proceeds, the mass differences carried by the connecting edges, the candidate structures that each path implies, and the fragmentation evidence by which the reasoning step selects among them. d) MS/MS are used to further reason for possible structural modifications between the query and neighborhood node. e) Additional information, such as prediction of structure with other tools, predicted formula for the parent mass etc. can be further included for all nodes in the propagation neighborhood.

The ego network alone can support inference - the probability of a match in the first layer is 8%, rising to over 30% when including approximate neighbor (suspect) matches(*19*), so it is already likely to contain spectral matches to structural relatives of the query. This expansion can be performed on either of two representations: the full global network, which retains all pairwise similarity edges and therefore recruits the densest possible set of second- and higher-order neighbors, or the MCN, whose spanning-tree structure restricts propagation to the strongest-edge backbone and yields a sparser, more conservative neighborhood. In either case the recruited neighbors may span multiple global molecular communities (Figure 1). The union of the ego network and this expansion, which we term the “propagation neighborhood”, is the information on which all subsequent inference is performed.

Each element of the propagation neighborhood constrains possibilities for the structure of the query. Edges carry mass differences that constrain the chemical relationship between the nodes they connect, annotated nodes suggest structures, and the query contributes its own precursor mass and MS/MS spectrum that can be compared with MS/MS spectra of nodes within the propagation neighborhood. Candidate structures are generated by propagating the structure of each annotated neighbor along the connecting path and applying the modifications implied by the mass differences encountered (Figure 1). A well-annotated neighborhood supports many such paths at once. Because the paths may be mutually inconsistent and the annotations from which they originate vary in reliability, the candidates are then resolved by reasoning over edge quality (cosine similarity) and fragmentation evidence, which discriminates among isomeric and isobaric candidates that mass differences alone cannot separate. A single consensus structure is then returned for the query.

This reasoning can be carried out by a trained mass spectrometrist, but it can also be performed by agentic AI, which has been shown capable of resolving molecular structures from mass spectrometry data(*20*) and is in general increasingly leveraged in metabolomics(*21*). Here we used frontier reasoning models, Claude Opus 4.8 (Anthropic), Grok 4.5 (xAI) and GTP-6 Sol (OpenAI), run independently on identical propagation neighborhoods (Methods).

### Annotation accuracy across authentic standards

DEEPNET-LLM was evaluated on 900 authentic standards of biologically-relevant molecules acquired by hydrophilic interaction (HILIC) and reversed-phase (RP) chromatography in positive and negative ion modes, comprising 2,249 scoreable spectra. The evaluation was fully blind: the identity of each standard was withheld from the models, and predictions were compared with the known structures only after all predictions had been returned. A prediction was scored as correct only if its full InChIKey, which encodes stereochemical configuration in addition to atomic connectivity, matched that of the standard.

We chose this collection of standards as the primary benchmark because no model could have encountered it. The standards are internal reference material that has never been deposited in a public domain, so the reported accuracy cannot be attributed to memorization or data leakage. Of note, public benchmarks offer no such guarantee: CASMI answers have been published, and MassSpecGym is assembled from public spectral libraries. MassSpecGym is therefore reported as a secondary benchmark, for comparison with other methods.

Reasoning with best performing model using DEEPNET-LLM assigned the top 1 correct structure to 84.7% of the 2,249 spectra (95% confidence interval (CI) 83.2–86.1%; Table 1, Figure 2, Supplementary Table 1); given this accuracy, only the top-ranked prediction was evaluated, without the top 10 and Tanimoto similarity reported conventionally. The two sources of evidence available to the reasoning step contributed unequally. Reasoning over the network alone, using only the information for immediate annotated neighbors and the mass differences on connecting edges to second-order neighbors, resolved the query with approximately 80% accuracy. Adding MCN hops to the propagation neighborhood beyond n=2 used here did not appear to improve accuracy further. Without the molecular network, using the MS/MS spectrum alone, no structure was predicted correctly (0%). Therefore, most of the structural determination constraint is ostensibly supplied by molecular context rather than by the spectrum of the query. This also corroborates the presumed absence of data leakage. Adding the MS/MS spectrum to the network raised accuracy in the top-ranked structure to above 90%, because fragmentation may further discriminate among isomeric and isobaric candidates that precursor mass differences leave degenerate.

**Figure 2.**
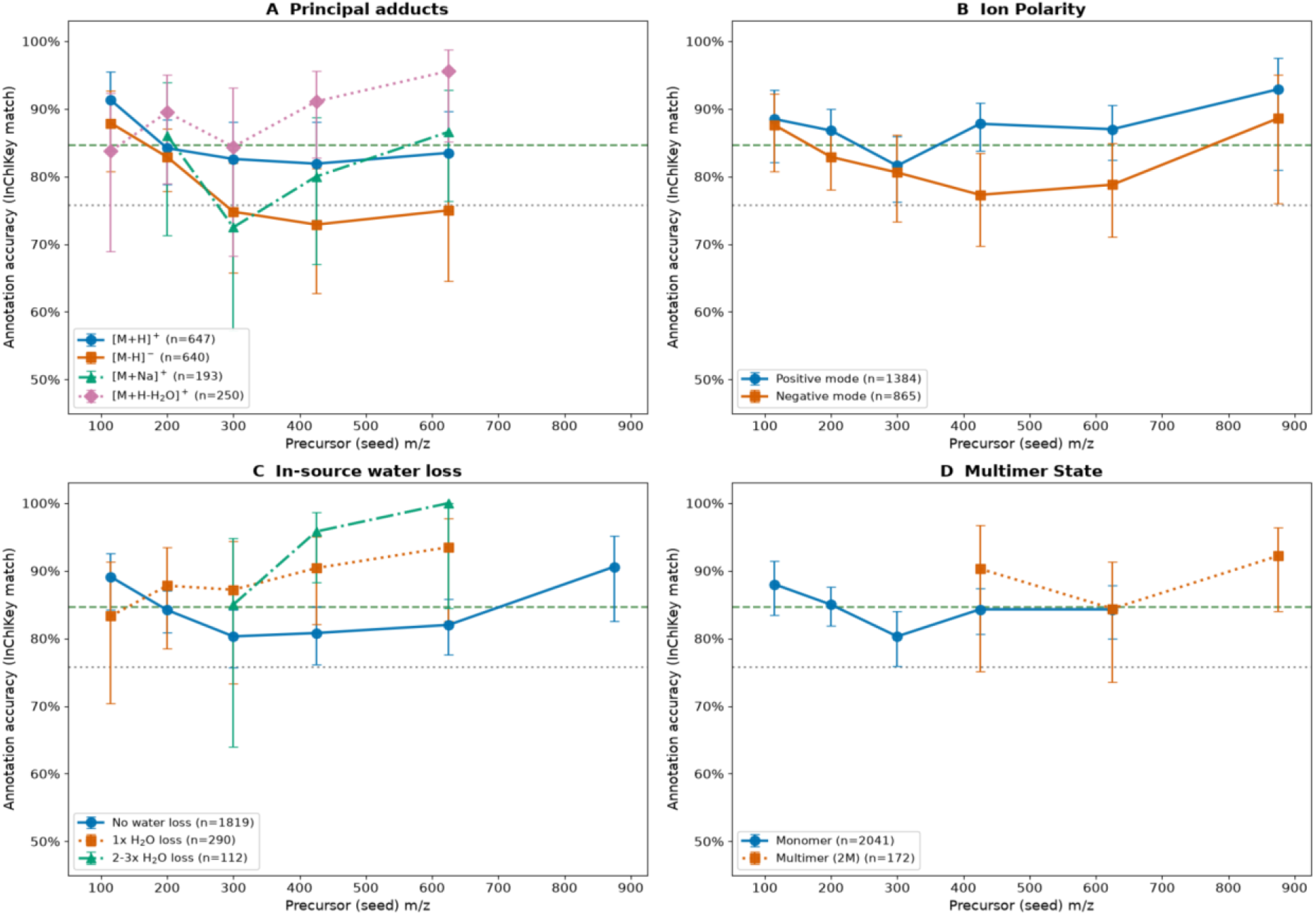
The prediction accuracy of DEEPNET-LLM vs. parent *m/z* of the query as a function of acquisition conditions. Predicted spectra are binned into 100 Da bins across the mass range, average accuracy is calculated. a) Ionoforms. b) Ion modes. c) Water loss. d) Multimers.

**Table 1.** Annotation accuracy across acquisition scenarios. Exact full-InChIKey (Top-1) match against 900 authentic standards, n = 2,249 scoreable spectra. The complete scenario breakdown is given in Table S1.

| Scenario | Accuracy | n |
|---|---|---|
| **All scoreable predictions** | **84.7%** | **2,249** |
| *Ion polarity* |  |  |
| Positive | 86.5% | 1,384 |
| Negative | 82.0% | 865 |
| *Principal adducts* |  |  |
| [M+H]<sup>+</sup> | 84.3% | 650 |
| [M−H]<sup>−</sup> | 80.1% | 643 |
| [M+H−H<sub>2</sub>O]<sup>+</sup> | 89.6% | 250 |
| [M+Na]<sup>+</sup> | 81.5% | 195 |
| *Multimer state* |  |  |
| Monomer | 84.2% | 2,050 |
| Multimer (2M) | 90.5% | 199 |
| *In-source water loss* |  |  |
| None | 83.3% | 1,819 |
| 1 × H<sub>2</sub>O | 88.6% | 290 |
| 2–3 × H<sub>2</sub>O | 95.0% | 140 |

Of the 343 incorrect predictions, 270 (79%) had the correct molecular formula but the wrong structure: propagation had reached the correct region of chemical space, and the reasoning step then selected the wrong isomer. Only 21% of all errors had an incorrect formula, the signature of a propagation or mass-assignment failure stemming from query placement into an incorrect neighborhood and/or insufficient information in the propagation neighborhood. Most errors were also minor: 70% of incorrect predictions were structurally close to the true compound, with a mean Tanimoto similarity of 0.66. The residual difficulty is therefore discrimination among isomers, not localization of the query in chemical space.

DEEPNET-LLM also exposed errors in reference data. As an example, the spectrum at *m/z* 182.0811 deposited as tyrosine was scored as a miss when both models returned para-tyrosine with high confidence. The structure, InChIKey and CAS number stored under that name were those of meta-tyrosine (3-hydroxyphenylalanine, CAS 587-33-7). Because DEEPNET-LLM reasons across an entire neighborhood rather than against a single library record, an annotation whose name, structure and network position are mutually inconsistent can be flagged.

The confidence reported with each prediction was informative. Accuracy rose monotonically with confidence, from roughly 70% at intermediate values to 87.5% for the 56% of predictions issued with a confidence of 0.9 or above. The reliability of an individual annotation can therefore be read from its score, and high-confidence predictions can be identified before any experimental confirmation.

### Accuracy is decoupled from the size of chemical space

The difficulty of structural annotation usually scales with the number of possible structures, which grows steeply with mass. The number of formulas consistent with an accurate mass of CHNOPS formulas under standard chemical constraints rises with m/z, and the number of structures consistent with those formulas rises far faster. At the upper end of the mass range, permissible structures exceed those actually observed by some 39 orders of magnitude (Figure 3A). Within a ±5 ppm window, the candidate set facing a method that searches chemical space at large grows by 30 orders of magnitude between the lowest and highest precursor masses examined (Figure 3B).

**Figure 3.**
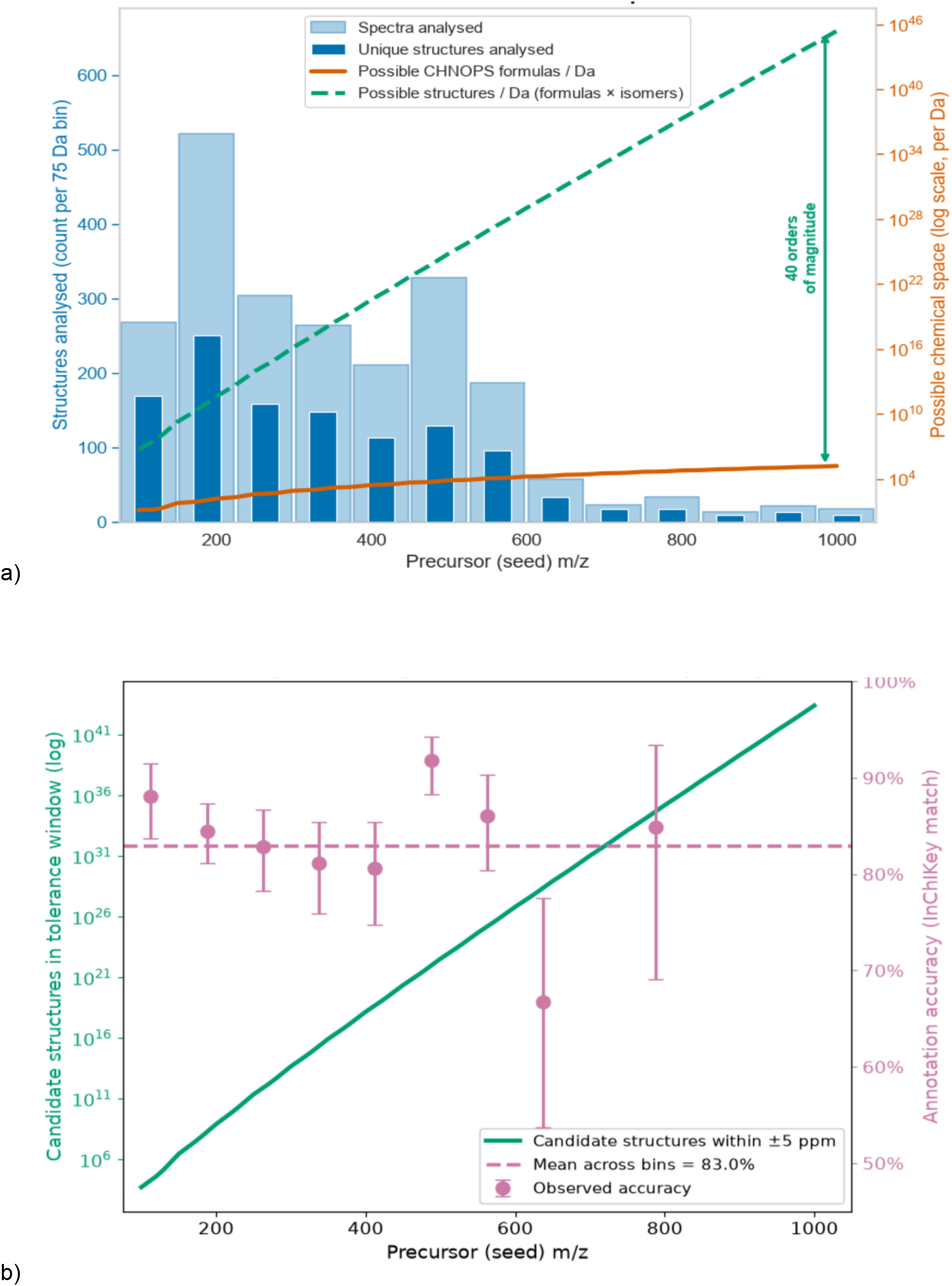
The prediction accuracy of DEEPNET-LLM is decoupled from the size of chemical space. a) Number of predicted structures (blue, left scale) across the m/z range vs number of possible formulas (orange, right scale) and structures (green, right scale). b) Prediction accuracy for compounds for 100 Da *m/z* bins, average accuracy is calculated (pink, right scale) across *m/z* range and the number of possible structures (green, left scale).

DEEPNET-LLM accuracy, by contrast, remained close to its mean of ∼84% across the same interval (Figure 3B). This decoupling is the central observation of this work. A method that draws candidates from chemical space must degrade in accuracy as that space expands, whereas DEEPNET-LLM that draws them from a molecular neighborhood is limited only by the density of information in the neighborhood, not by the size of the space.

### Consistency across acquisition conditions

Accuracy was found to be consistent across acquisition conditions (Figure 3 a, b, Table 1). It was effectively independent of ion polarity (86.5% positive, 82.0% negative), although the two polarities involve different ionization chemistry and produce dissimilar fragmentation, indicating that the constraint supplied by the propagation neighborhood is sufficient for both ionization modes. Accuracy was further comparable for all principal adducts, including protonated, deprotonated, sodiated and water-loss species (no ammonium and potassium adducts in this test set).

Multimers were annotated as accurately as monomers, and spectra showing in-source water loss at least as accurately as those without (Figure 3c, d; Table 1). Accuracy was consistent across *m/z* in every condition examined, in agreement with its independence from the size of the candidate space.

These data also quantify how much of the feature count arises from ionoforms and in-source fragments (ISFs). Protonated and deprotonated ions accounted for 57.5% of features; the remaining 42.5% were in-source water-loss forms (18.9%), adducts (14.2%) and multimers (8.8%) of the same parents. Grouped by compound of origin, the 2,249 spectra collapsed into 798 distinct metabolites, so that 35.5% of features represented distinct chemistry. Redundancy was greater in positive mode, where 48.1% of features were unique, than in negative mode (62.4%), consistent with the richer adduct and water-loss chemistry of positive ionization.

### Performance on MassSpecGym

DEEPNET-LLM was also evaluated on MassSpecGym(*12*), for comparison with published methods. Following the convention of the benchmark and of the *de novo* structure prediction literature, stereochemistry was excluded to provide direct comparability (a prediction was scored as correct when the first block of its InChIKey, which encodes atomic connectivity, matched that of the reference structure). This criterion is less stringent than the full-InChIKey match applied to the authentic standards, which, rescored on connectivity alone, reach 89.9% (Supplemental Table 1).

The evaluation set comprised random 285 test-fold spectra (Methods). All inputs were blinded: the name, structure and database accession of each query were removed from every model-facing file, reference structures were held separately until scoring, and each spectrum was reasoned over individually, without pre-training, fine-tuning or benchmark-specific adaptation. In the library-assisted configuration, the spectra of neighbors, but never of the query, were searched against the NIST 2023 MS/MS library, and the resulting matches were supplied to the model.

With neighbor-only library search, Claude Opus assigned the correct structure to 55 of 64 spectra (85.9%; 95% CI 75.4–92.4%), 52 of which (81.2%) also matched the reference structure exactly (Table 2). This run was terminated after 64 spectra because of its pre-set token cost. Without library search, Grok 4.5 annotated 102 of the 285 spectra correctly (35.8%; 95% CI 30.4-41.5%). Most of its shortfall reflected abstention rather than error: Grok returned no structure for 128 spectra, and 65.0% of the 157 structures it did return were correct. With the same inputs as Claude Opus, a locally run open-weight model, Qwen3-14B, reached 13.8% (12 of 87 spectra; partial run), so accuracy depends on the reasoning capability of the model. A chemistry language model given the spectrum without a propagation neighborhood (ChemDFM-R) reached 1.1%, mostly by proposing structures inconsistent with the precursor mass. The best configuration exceeds the top-1 accuracy of leakage-controlled *de novo* methods by almost two orders of magnitude (Table 2).

**Table 2.** Annotation accuracy on MassSpecGym. Top-1 accuracy is scored on the first block of the InChIKey (atomic connectivity, stereochemistry excluded), following the convention of the benchmark; exact match denotes canonical SMILES identity. Evaluation set: 285 test-fold spectra (18 unique structures). 95% CIs are Wilson intervals. *Partial run (Claude Opus stopped for token cost; Qwen3-14B stopped at runtime limit). †Grok returned no structure for 128 spectra; accuracy on returned structures, 65.0% (102 of 157). ‡Reference values from MassSpecGym in the Wild, 2026(*13*): range of published top-1 accuracies, with best top-10 accuracy in parentheses.

| Configuration | Spectra scored | Top-1 accuracy (n correct) | 95% CI | Exact match (n) |
|---|---|---|---|---|
| *Propagation neighborhood + neighbor-only NIST search* |  |  |  |  |
| Claude Opus 4.8 (Anthropic)* | 64 | 85.9% (55) | 75.4–92.4% | 81.2% (52) |
| Qwen3-14B (open-weight, local)* | 87 | 13.8% (12) | 8.1–22.6% | 13.8% (12) |
| *Propagation neighborhood, no library search* |  |  |  |  |
| Grok 4.5 (xAI)† | 285 | 35.8% (102) | 30.4–41.5% | 31.6% (90) |
| *No propagation neighborhood* |  |  |  |  |
| ChemDFM-R | 285 | 1.1% (3) | 0.4–3.0% | 0.7% (2) |
| *Published methods, leakage-controlled split‡* |  |  |  |  |
| De novo generation, current state of the art | — | 0–2% (top-10: 4.1%) | — | — |

These results carry two caveats. First, the 285 spectra cover 1.6% of the 17,556-spectrum test fold and represent only 18 unique structures, so spectrum-level accuracy is dominated by replicate spectra of a few molecules. Second, because MassSpecGym spectra derive from public libraries, data leakage is possible in theory. The training dataset has not been used, and the model did not encounter it prior to prediction, but this does not eliminate the data leakage possibility entirely, as the predicted information may have been encountered during the frontier models’ training.

### Mapping of molecular space

Because candidate structures are drawn from the propagation neighborhood, the quality of prediction ostensibly depends on how well the surrounding molecular space is populated with high-quality annotations. A neighborhood in which annotations are sparse, so the query node is connected to annotated nodes only through distant or low-similarity edges, supports fewer, longer and less reliable propagation paths, potentially impairing prediction accuracy. This complicates annotation but does not altogether prevent it, as structures can still be propagated from suspect-level annotations (*19*) or over longer distances. The “darkest” parts of the global molecular network are expected to be the least amenable to DEEPNET-LLM. To determine how much of the mapped metabolome falls into such regions of the global network, we computed three statistics for every node of the global MCN(*17*): the number of hops to the nearest annotated node, the fraction of annotated spectra among the k = 25 first-order matches of the ego layer, and the distribution of edge cosine similarity as a function of distance from the query (Table 3; Methods).

**Table 3.**
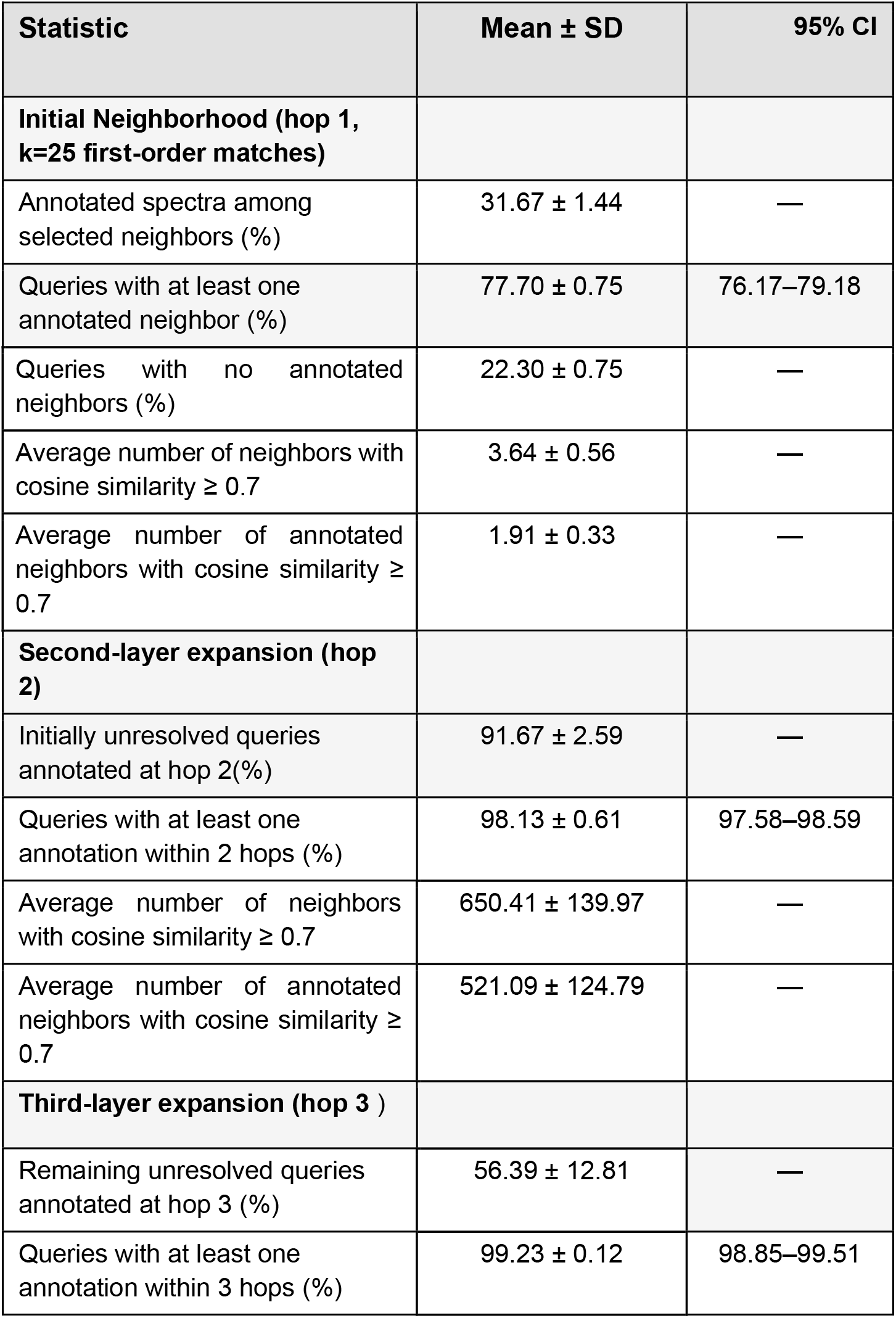
Annotation statistics of the global MCN(*17*). Results are based on 1,000 query spectra independently sampled with replacement for each of three random seeds. Annotation availability was assessed among the 25 highest cosine first-order neighbors and following expansion to the second and third hops; both GNPS library and nearest annotated neighbor(*19*) matches are counted. High similarity neighbors at hops 1 and 2 were identified using an edge cosine similarity threshold of ≥ 0.7. Values are reported as mean ± SD across seeds. The 95% confidence intervals (CIs) for annotation coverage were calculated using the exact binomial method.

**Table 4.** predictions for the synthetic sugars (negative ion mode). Formula and adduct were correct in all 12 predictions; constitution (InChIKey connectivity) matched in none. MCES: myopic MCES, threshold 15. Tanimoto: ECFP4 (radius 2, 2048 bits). *Compared with the open-chain form of the true compound, because the prediction was drawn open-chain; against the pyranose form MCES is 4.

| Parent Mass | SMILES | Correct Structure | Predicted Structure by SIRIUS | Predicted Structure by DEEPNET-LLM |
|---|---|---|---|---|
| 193.07197 | CC(O)[C@H]1O[C@H](O)[C@H](O)[C@@H](O)[C@@H]1O |  |  |  |
| 255.08748 | OC(c1ccccc1)C1OC(O)C(O)C(O)C1O |  |  |  |
| 207.0878 | CC(C)(O)[C@H]1O[C@H](O)[C@H](O)[C@@H](O)[C@@H]1O |  |  |  |

These statistics indicate that, on average, every query node has approximately 2 annotated neighbors with cosine similarity ≥ 0.7 in the ego layer, and that essentially no nodes lay farther than three hops from an annotated one. Thus, at the current rate of annotation, the global molecular network appears to already contain enough structural information to anchor propagation for nearly all of the mapped metabolome, supporting the general applicability of the method. Queries with no annotation in the ego layer (22.30% of all features) may still be resolved by expansion to the second layer if/when they are encountered. However, in such cases, annotation is indeed impaired: manual examination of queries with incorrect annotations of test data confirmed that the predicted structures were often drawn from poorly-populated neighborhoods. Consequently, the most straightforward way to further improve prediction accuracy is to populate more of the global network with annotations, through continued deposition of annotated reference spectra and community curation.

### Limits of the prediction

The evidence presented above indicates that DEEPNET-LLM draws most of its structural information from the propagation neighborhood rather than from the query spectrum alone, a unique feature of the methodology. Correspondingly, its behavior, and therefore its failure modes, should be predictable. Structures that meet the following two conditions are expected to be incorrectly annotated: the exact structure is 1) not resolvable from MS/MS, and 2) contains structural features that cannot be anticipated, in particular in cases where both the annotated neighbors and the priors of the models point toward a different, more common structure. To verify this behavior, we designed such a test set using synthetic sugars.

We synthesized hexopyranose analogues in which the C6 hydroxymethyl group is replaced by an unusual carbon substituent: CH(OH)CH_3_ (C_7_H_14_O_6_), C(OH)(CH_3_)_2_ (C_8_H_16_O_6_) or CH(OH)Ph (C_12_H_16_O_6_) (Methods). Each compound shares its molecular formula with a common, biologically plausible isomer: the methyl, dimethyl and phenyl O-glycosides of a hexose, which are known or potential natural products of *Streptomyces* that inhibit carbon dioxide fixation by *Scenedesmus obliquus*. We acquired negative-ion MS/MS spectra for all three ([M−H]^−^ at *m/z* 193.0720, *m/z* 207.0878 and *m/z* 255.0875).

The spectra contained information that can be used for structural hypothesis generation, with 10-15 fragment ions above 5% relative intensity and precursor fractions of 0-6.4%, but exact isomers would not be resolvable from MS/MS spectra alone. The MS/MS spectra were dominated by the generic carbohydrate cross-ring ions *m/z* 59.014 (C_2_H_3_O_2_, base peak in all three), *m/z* 89.025 (C_3_H_5_O_3_) and *m/z* 71.014 (C_3_H_3_O_2_). These ions report on the glycan core, not on the position of the modification. The few ions that carry the substituent mass (*m/z* 103.041 and *m/z* 73.030; *m/z* 147.067 and *m/z* 117.056; 135.046) are equally consistent with substitution at the anomeric oxygen.

We found that this query has indeed returned an incorrect structure. Without additional context, DEEPNET-LLM returned the common O-glycoside isomer for every compound, in all three evidence settings (propagation neighborhood only; with SIRIUS formulas; with SIRIUS formulas and FRIGID/ICEBERG candidates) (Table 1). The molecular formula and adduct were correct in all nine predictions; SIRIUS rank-1 formulas agreed (scores 43.7-47.8; 77-90% of MS/MS spectral intensity explained), but the exact structure was incorrect in all nine. All three settings converged on the same connectivity for each spectrum, and stated confidence was moderate to high (0.40-0.70). The predictions were structurally close, where each differs from the true compound by relocating a single substituent (MCES 2-4), which ECFP4 similarity scores as distant (Tanimoto 0.25–0.33).

We then repeated the annotation with an identical query and an additional prompt stating that the compounds are synthetic, so that biological plausibility and library frequency should not be used as priors. With this context, DEEPNET-LLM abandoned the O-glycoside for two of the three compounds. For *m/z* 193.072 it proposed a 4-deoxyheptopyranose and for *m/z* 207.088 an open-chain 6,8-dideoxyoctose: chain-extended reducing sugars, the correct structural class, each within MCES 2 of the true compound. Neither matched the exact substitution pattern; the model placed the deoxy position or branching differently. For *m/z* 255.088 the model proposed a different glycoside, 2-(3,4-dihydroxyphenyl)ethyl tetrofuranoside, which is further from the true structure than the original call (MCES 6). Stated confidence decreased to 0.30-0.40, consistent with the model recognising that the spectra do not localise the modification.

This outlines limitations of DEEPNET-LLM, but also supports the conclusion that the method does not fail arbitrarily. It returns the most plausible isomer consistent with the spectrum and its neighborhood, with the correct molecular formula and adduct and a connectivity close to the true one. Supplying context that removes the biological prior moved two of the three predictions to the correct structural class and lowered stated confidence accordingly, showing that the predictions respond to the evidence and context provided. This further defines how the method’s results should be interpreted. For features from biological samples, where the biological prior is expected to hold, the propagated structure should be the most likely answer. For compounds that may lie outside familiar biochemistry, such as synthetic or xenobiotic chemistry, predictions should be treated as class-level hypotheses whenever the MS/MS does not localize the modification, and any relevant knowledge about the sample should be supplied to the models.

### Example of molecular discovery

We have applied DEEPNET-LLM to predict unknown spectra in experimental data. Gut samples from humanized mouse model and a gut bioreactor have been examined by first constructing a molecular community network from the data. A large community of features that predominantly contains annotations consistent with various bile acids (Figure 4) was further investigated in greater detail, as a stream of recent discoveries, starting from microbial conjugation(*22*), has significantly increased interest in this molecular family. In one example, we have identified a feature with a parent *m/z* that did not match any known conjugate possibilities, including various amino acids, but, assuming a deoxycholic acid (DCA) cholate core, matched conjugation of a moiety with the mass of serotonin. This structure has no immediate annotated neighbors and thus conventional annotation propagation was not possible. We applied DEEPNET-LLM for the structure prediction, with no additional information supplied to the models. All three reasoning models (Claude, Grok and Gemini) returned N-cholyltryptamine, 3α,7α,12α-trihydroxy-N-[2-(1H-indol-3-yl)ethyl]-5β-cholan-24-amide, agreeing on connectivity at confidences of 0.76, 0.60 and 0.77, with two of the three also assigning the 3α,7α,12α configuration (Figure 4). The exact assignment rests on the tryptamine-based MS/MS spectral ions at *m/z* 144.080 and *m/z* 161.106, in place of the *m/z* 160.076 and *m/z* 177.102 that serotonin would yield, and on the trihydroxy steroid losses at *m/z* 337.250 and *m/z* 319.240, in place of the 339 and 321 of a dihydroxy bile acid, manually confirmed by an expert mass spectrometrist. A synthetic standard of tryptamine conjugated to cholic acid reproduced the full fragment series of the unknown, as well as the retention time, confirming the structural assignment at Level 1(*23*).

**Figure 4.**
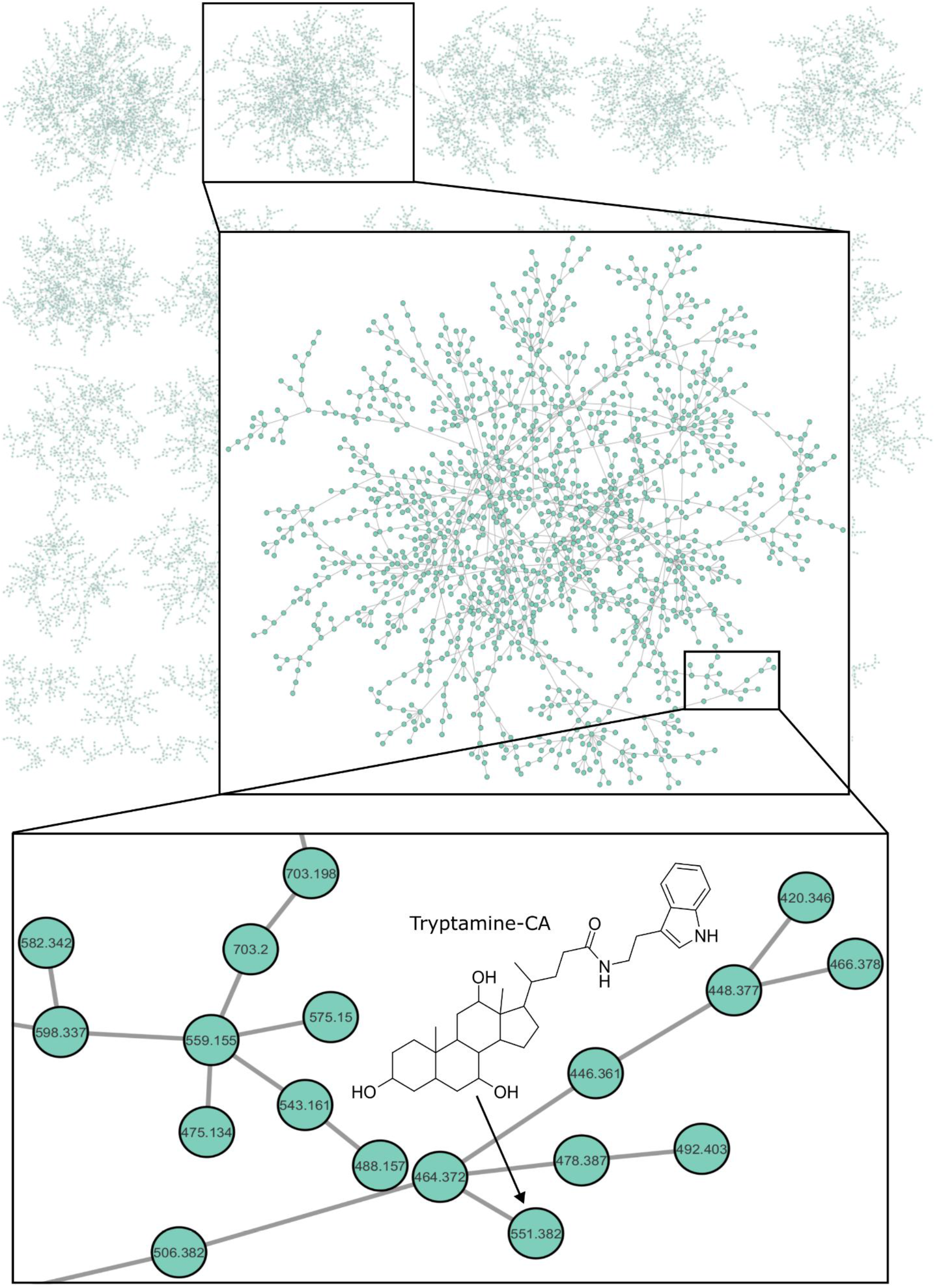
Discovery of tryptamine-CA conjugate. The full community network of samples from humanized mouse model and gut bioreactor samples with the highlighted community that contains multiple bile acids annotations. The feature of interest is therefore presumed to be a bile acid-related compound.

This identification is notable as the N-cholyltryptamine is, to our knowledge, a previously unreported metabolite. It extends the rapidly growing family of microbially conjugated bile acids and links two major axes of host-microbe chemical communication, bile acid signaling and microbial tryptophan metabolism. This finding also demonstrates that the path from an unannotated feature to a Level 1 identification now may require the synthesis of just a single standard, rather than the synthesis and testing of multiple candidate structures.

## Discussion

The structure prediction approach demonstrated here, *De novo* Ego network-Enabled Propagation and Neighborhood Extrapolation of Topology using Large Language Models (DEEPNET-LLM), is based on organizing metabolome by constructing global molecular community network (MCN) to capture structural patterns and leverage similarities with molecular neighbors in a “propagation neighborhood”. MCN was able to overcome limitations of conventional molecular networking, which uses a single similarity threshold, and due to greatly varying patterns in MS/MS spectral features across molecular families, both over-connects some nodes and disconnects others. This results in artificially fractured and distorted clustering patterns. Furthermore, in conventional networks more than half of all features are disconnected (so-called singletons), and many of the small fractured clusters carry no annotation from which to propagate. In contrast, MCN connects more than 95% of features, capturing natural patterns in spectral similarities, and provides an annotated neighbor for approximately three million previously isolated features across GNPS(*17*).

At the current level of annotation, a global neighborhood grown around a query is already likely to contain annotated structures in its immediate vicinity, and for all queries, not further than 3 hops away (Table 3). The annotation rate observed here against the current GNPS library, including approximate structures from the “Suspects” list(*19*), is approximately 30%, and our results show that this density of information is sufficient to facilitate correct top-1 prediction in more than 80% of cases. The information within propagation neighborhoods could be further enhanced via additional library searches, formula and structure prediction with other tools (Methods). We found that supplementing networks with the results of SIRIUS/CSI FingerID, for example, does discernably improve prediction accuracy. Earlier automated propagation assigned only compound classes across subnetworks (*11*); DEEPNET-LLM returns an exact structure for each query.

The use of agentic AI in the proposed approach, as described and demonstrated here, is not strictly necessary, as the same reasoning over the propagation neighborhood can, in principle, be conducted manually. However, AI substantially accelerates the process thus enabling far greater throughput necessary for large-scale data analysis. Because the reasoning and provenance behind each proposed annotation could be manually validated, the AI use in this context is not a “black box”. A chemist/mass spectrometrist can also construct prompts in a way to safeguard against certain choices or otherwise influence reasoning based on available information. Also, although the possibility of AI “hallucination” is generally considered a problem, in the present context it enables proposing novel structures. Human validation of the proposed structures is still essential, however.

The presented results support the central premise of molecular networking, that spectral similarity reflects structural similarity. Manual examination confirms that DEEPNET-LLM generally places query spectra in their correct molecular neighborhoods (the node is surrounded by neighbors with annotations that are generally consistent with that of the query, when it is known), which is further evidenced by the downstream high prediction accuracy. In the absence of data leakage, this is only achievable if prediction of the query spectrum is drawn from similar structures in its neighborhood. The fact that the complete structure, including stereochemistry, can be predicted at all is more surprising, because MS/MS fragmentation does not uniquely define configuration. A likely explanation is that because AI is probabilistic, and not deterministic, it predicts the likely stereochemistry that was encountered in LLM’s training, which would be that generally realized in living systems. The biological stereochemistry is a small, non-uniformly distributed subset of what is chemically permissible, and this information is inherited from annotated neighbors by annotation propagation. The observed errors support this interpretation, as they tend to concentrate where biological configuration is itself variable, notably among bile acids (Supplemental Table 1), whose many stereoisomers occur across biology and cluster together without a dominant form for the method to select. The errors are rare where configuration is conserved: for example, amino acid stereochemistry, near-uniform across living systems, was always predicted correctly. DEEPNET-LLM is therefore accurate to the degree that biology is regular, consistent with the observation that most structural information is drawn from the network rather than from the individual spectrum. A case of modified glycans described in Results that was constructed deliberately to make DEEPNET-LLM fail, further illustrates this. Because propagation draws on what biologically plausible, and the models themselves are weighted toward structures well represented in training and in reference databases, DEEPNET-LLM can be misled by a structure that departs from the expected isomer in a way the MS/MS fragmentation does not capture, and when no further evidence is supplied to indicate that it should. Annotations of compounds that may lie outside biologically “familiar” chemistry, should therefore be treated as class-level hypotheses if MS/MS cannot localise a modification.

The accuracies reported here are expected to be a lower bound. The models were used as supplied, with a fixed prompt that was not tuned, and without task-specific training or selection among outputs, because such interventions, while they may raise performance, risk providing “cheats” leading to overfitting and reduced generality. Accuracy should naturally improve as the reference global network grows, since every added MS/MS spectra extends the mapped chemical space and every new annotation adds context for future queries. Because accuracy is limited by annotation density rather than by the size of chemical space, Table 3 also identifies that there exist regions of the network that are more “dark” than others and addition of structural references for those regions of the global network could further improve prediction accuracy. However, at the accuracy already achieved, confirming a novel identification in ∼85% of cases can be reduced to testing a single, high-probability candidate. Importantly, DEEPNET-LLM is orthogonal to all other existing *in silico* tools as it is based on a different principle. It is possible to combine predictions from other methods across propagation neighborhoods to potentially improve reasoning.

The practical consequence of high-accuracy structure prediction is that molecular discovery can be greatly accelerated, as illustrated by the discovery of N-cholyltryptamine. The feature was found within a bile acid community of the MCN constructed from the study data, which, at the dataset level, only suggests an association with the bile acid molecular family. DEEPNET-LLM resolved the complete structure from the data, and a single synthetic standard was sufficient to confirm it at Level 1. Amidation of bile acids beyond the host glycine and taurine conjugates has been recognized only recently(*22*), and of amines that are not amino acids only the GABA and tyramine conjugates have been described, with no function assigned to either(*24*, *25*). N-cholyltryptamine discovery extends this class to tryptamine, a bacterial decarboxylation product of tryptophan that acts on the colonic epithelium through the 5-HT₄ receptor(*26*, *27*). The conjugate thus joins two of the principal chemical languages through which the gut microbiota communicates with the host: bile acids, which signal through receptors such as FXR and TGR5 (*28*, *29*), and microbial tryptophan metabolites, which regulate gut motility, secretion and immune function (*30*, *31*). Both halves of the molecule carry established signaling roles, and the conjugate itself may possess a signaling function distinct from either, for example engaging bile acid and serotonergic receptors at once.

The existence of N-cholyltryptamine also could be considered in the context of the fate of tryptamine in the gut. Free tryptamine is metabolized by monoamine oxidase (MAO) to indole-3-acetic acid (IAA)(*32*). Amidation of the primary amine would be expected to protect tryptamine from oxidative deamination, so conjugation could act as a reservoir that prolongs its availability, a sink that removes it from 5-HT₄ signaling, or a route by which it enters enterohepatic circulation together with the bile acid. Because bacterial bile salt hydrolases can act as acyltransferases that conjugate bile acids to amines (*33*), the conjugate may represent a further point at which microbial metabolism couples the bile acid pool to tryptophan-derived signaling. Which of these roles, if any, it plays is an open question, but one that can now be addressed directly, since the structure is established and the compound is synthetically accessible. More broadly, the discovery of a previously undescribed metabolite at the intersection of two of the most intensively studied pathways of the gut metabolome suggests how much remains undescribed, and how much becomes reachable once structure prediction can be automated and applied routinely.

Finally, the ability to propose high-accuracy annotations, including those of ionoforms and ISFs, allows the detected metabolome to be separated into distinct chemistry and redundant variants (adducts, in-source fragments and multimers) of common parents. The compounds examined as described in the “Annotation accuracy across authentic standards” section of Results were chosen for their importance across biology and span a wide chemical range, so the patterns observed for them should be, at least approximately, representative of patterns in biological samples. Across all MS/MS spectra used in testing, 35.5% of features represented distinct chemistry, which is approximately equal to the fraction correctly predicted by DEEPNET-LLM (35.2%). Assuming similar ratios in experimental data, across 8.4 million indexed GNPS spectra, of which roughly 92% are currently unannotated, this implies two to three million previously undescribed metabolites that are not ISFs. Noting that a curated set cannot reproduce the redundancy of an arbitrary sample, it is important to point out that the proportion of ISFs is substantially below the high artifact fractions estimated previously(*34*). Importantly, such estimates can now be directly made from the data via structural assignment of individual features, because the structures of in-source fragments and ionoforms are themselves predictable with high accuracy.

Taken together, the accuracy of DEEPNET-LLM, its independence from molecular size and its stability across acquisition conditions indicate that annotation of previously unreachable features may become routine. The dark matter of metabolomics, long treated as an intrinsic limit of the measurement, should instead be considered as a consequence of attempting to interpret each spectrum in isolation. It recedes once each spectrum is read in the context of the molecules to which it is related.

DEEPNET-LLM also has limitations, in addition to those noted above. DEEPNET-LLM depends on annotated neighbors, and although sparse annotation complicates rather than prevents propagation, the 22.3% of the global network with no immediate annotated neighbors (Table 3) is the least amenable to the approach, generally leading to lower prediction accuracy. The propagated information inherits errors in reference annotations, although reasoning across a neighborhood can expose some of them. Stereochemistry is recovered only to the extent that it is conserved in biology and any stereochemical assignments must be treated with skepticism. Finally, reasoning with frontier models is currently computationally costly, which limits evaluation at repository scale (on MassSpecGym, a single model consumed approximately 8 million tokens for 64 spectra).

## Materials and Methods

### Construction of the propagation neighborhood

The reference index against which every query is placed comprises all publicly available GNPS MS/MS spectra following clustering to remove redundancy, amounting to 8.4 million spectra in the version used here. Indexing and search were performed with HNSW-MS (*18*), which implements hierarchical navigable small world graph indexing natively for mass spectral similarity. The implementation operates directly on raw spectra, each represented as a sparse collection of mass-to-charge values paired with intensities, without binning, preprocessing or transformation into fixed-dimensional embeddings, so that the similarity scores returned by the index are identical to those obtained by exhaustive pairwise comparison and the retrieved neighborhoods are fully reproducible. Similarity is computed by matching peaks within an *m/z* tolerance window of 0.01 Da, followed by greedy assignment of the matched peaks and cosine scoring of the matched intensity vectors.

The index is a multi-layer graph in which each spectrum is assigned a maximum layer drawn from an exponential distribution governed by *M*, the target number of bidirectional connections per node, and in which insertion proceeds by greedy search from the graph entry point to identify candidate neighbors followed by the establishment of bidirectional edges to the *M* closest candidates at each layer. Because cosine spectral similarity does not satisfy the triangle inequality, the pruning heuristics on which most approximate nearest neighbor methods rely are not available, and connections are therefore allowed to accumulate without aggressive pruning, which preserves recall in this non-metric space at a modest cost in memory (*18*). The index used throughout this work was built with *M* = 16, efConstruction = 200 and queried at efSearch = 128, settings at which repository-scale search returns the correct best-matching spectrum for 95% to 98% of queries under the relaxed top-1 criterion. Index construction is a one-time operation, requiring approximately 12 hours on a single CPU core at these settings, after which the index is queried indefinitely and accepts incremental insertion of new spectra without reconstruction.

A query spectrum is placed by greedy routing from the sparse upper layers of the graph down to the dense base layer, which retrieves nearest neighbors in logarithmic rather than linear time and requires approximately 5.6 ms per query against the 8.4 million spectrum index. The *k* most similar spectra are returned, with *k* = 25 used throughout, and these first-order matches constitute the ego network of the query.

Each first-order match is then located within the global Molecular Community Network assembled across all public GNPS data (*17*), and the subnetwork is expanded outward by *n* hops, with *n* = 2 used here, recruiting second- and, if needed, higher-order MCN neighbors that may span multiple molecular communities. The union of the ego network and this expansion constitutes the propagation neighborhood. Every node in the propagation neighborhood carries any structural annotation deposited against it, together with the provenance and confidence level of that annotation and including approximate structures from the GNPS Suspects list(*19*), and every edge carries the precursor mass difference between the two spectra that it connects. The propagation neighborhood passed to the reasoning step therefore consists of the precursor mass and fragmentation spectrum of the query, the precursor masses and fragmentation spectra of the neighbors, the annotated structures as SMILES and InChIKey with their confidence levels, and the mass differences carried by the connecting edges.

The propagation neighborhood networks can be used as is for downstream prediction, or the information within can be further enhanced. Here, we tested multiple steps and compared their effect on the prediction accuracy with and without: additional library search using NIST 2023, molecular formula prediction using SIRIUS 4, and structure prediction with FRIGID(*35*).

### Reference standard library and spectral acquisition

#### Reference standards

The standards were biologically relevant metabolites(*1*) purchased from MetaSci (<u>spec.metasci.ca</u>), and their spectra were acquired as described previously(*17*). The resulting spectral collection has never been deposited in a public repository, published or otherwise released. To exclude instrument-to-instrument variation in MS/MS spectra while retaining the variation introduced by the mode of acquisition, all spectra were recorded on a single instrument using four methods: reversed-phase (RP) and hydrophilic interaction (HILIC) chromatography, each in positive and negative ionization mode. Standards were analyzed in batches of 50 by all four methods, so that each compound could be observed as several ionoforms. Each spectrum was labeled with the identity supplied by the vendor, and was retained only after that identity had been confirmed by search against existing spectral libraries, giving one curated library per method.

#### Chromatography

Separations were performed on a Vanquish LC system (Thermo Fisher Scientific) at a flow rate of 0.5 mL/min, with the column at 40 °C and the autosampler at 4 °C. RP separations used a CSH C18 column (Waters; 50 × 2.1 mm, 1.7 µm) eluted with a gradient of water (A) and acetonitrile (B), both containing 0.1% formic acid. HILIC separations used a BEH Amide column (Waters; 50 × 2.1 mm, 1.7 µm) eluted with a gradient of water containing 10 mM ammonium formate and 0.01% formic acid (A) and acetonitrile containing 10 mM ammonium formate and 0.1% formic acid (B)(*17*).

#### Mass spectrometry

Spectra were recorded on an Orbitrap Exploris 240 mass spectrometer (Thermo Fisher Scientific) with a heated electrospray ionization source, using data-dependent acquisition over a 10-min method. The spray voltage was 3.5 kV in positive mode and 2.5 kV in negative mode, and the ion transfer tube and vaporizer were held at 350 °C and 400 °C, respectively. Full scans were acquired at a resolving power of 30,000 and MS/MS scans at 15,000, and with the standard AGC target; in-source fragmentation was not applied. Precursor selection was guided by an inclusion list specifying the formula, expected adduct, *m/z*, polarity and retention-time window of each standard.

#### Data processing

Vendor files were converted to mzML and processed with MZmine 3(*36*). Features were detected at an intensity threshold of 5 × 10³ with peak widths of 2 s to 2 min, isotopic peaks were removed, and peak lists were aligned with tolerances of 5 ppm in *m/z* and 10 s in retention time.

#### Evaluation set and scoring

Of the 2,513 spectra acquired, 264 lacked a reference InChIKey and were excluded, leaving 2,249 scoreable spectra. The adduct, multimer and in-source water-loss state of each MS/MS spectrum was assigned via a manual examination by an expert mass spectrometrist. For structure prediction, the identity of each standard was withheld from the models, and predictions were compared with the reference structures only after all predictions had been returned. Predictions were scored by exact full InChIKey match (connectivity and stereochemistry), by first-block InChIKey match (connectivity only) and by Morgan fingerprint Tanimoto similarity computed with RDKit. Confidence intervals reported throughout are 95% Wilson score intervals.

### Synthetic reference compounds structure prediction

#### Hexopyranose synthesis

Each analog involves a substitution on the C6 position of either glucose or galactose. The 6-C-methylglucose and 6-C-phenylglucose were synthesized by protecting glucuronolactone with tert-butyldimethylsilyl (TBS) chloride in dimethylformamide. The protected molecule reacted with methyl lithium for the methyl analog and with phenyl lithium for the phenyl analog(*41*, *42*).The lactol intermediate underwent a diastereoselective reduction following the Felkin-Ahn model, then deprotected using 60% trifluoroacetic acid/water to yield the (S)-diastereomer. Removal of the tert-butyldimethylsilyl group prior to reduction of the lactol intermediate resulted in an inseparable mixture of diastereomeric triols. The mixture was reacted with acetone and camphorsulphonic acid to yield a mixture of diacetonides, which were separated via flash column chromatography and deprotected with 60% trifluoroacetic acid/water to yield the corresponding 6-C-methylglucose diastereomers. Contrary to this approach, the 6-C-phenyl triol diastereomers were readily separable. The 6-C-dimethylglucose synthesis was achieved by reacting the 5-O-silyl protected glucuronolactone starting material with methylmagnesium bromide. The diol product was then deprotected and recyclized using 60% acetic acid/water at 100 °C.

Three other molecules with the same substitutions were also synthesized for galactose at the C6 atom. All three analogs started with the oxidation of the OH group at C6 of diacetonegalactose. From here, the 6-C-dimethylgalactose and 6-C-phenylgalactose was synthesized using a series of grignard reactions and oxidation-addition reactions(*43–45*). For the 6-C-methylgalactose, it underwent Wittig olefination, epoxidation, separated the diastereomers, then LAH reduction to get the final product.

For carbohydrate analyses, samples were dissolved in 50:50 (v/v) acetonitrile:water to a final concentration of 5 mg/mL. Analyses were performed on a Vanquish UHPLC system coupled to an Orbitrap Exploris 480 mass spectrometer (Thermo Fisher Scientific). HILIC separations used an ACQUITY UPLC BEH Amide column (Waters; 100 × 2.1 mm, 1.7 µm) maintained at 40 °C, with the autosampler at 4 °C and an injection volume of 10 µL. Mobile phase A was water containing 0.1% formic acid and 10 mM ammonium formate, and mobile phase B was 95:5 (v/v) acetonitrile:water containing 0.1% formic acid and 10 mM ammonium formate (all solvents Optima LC–MS grade, Fisher Scientific). The flow rate was 0.4 mL/min, and the gradient was as follows: 100% B was held for 1.0 min, then decreased linearly to 70% B at 6.7 min, to 40% B at 7.0 min, and to 35% B at 8.0 min, where it was held until 9.0 min. The mobile phase was then returned to 100% B at 9.1 min and held until 10.5 min for column re-equilibration, giving a total run time of 10.5 min.

Mass spectrometry was performed in negative ionization mode over a mass range of m/z 75–1125. Source parameters included a spray voltage of 3,500 V, sheath gas pressure of 35 psi, auxiliary gas pressure of 10 psi, ion transfer tube temperature of 350°C, S-lens RF level of 50 and auxiliary gas heater temperature of 400°C. External mass calibration was performed prior to data acquisition using Pierce LTQ Velos ESI negative Ion Calibration Solution and mass accuracy was within 1 ppm. Data was acquired using data-dependent MS/MS mode with MS1 scans collected at 30,000 resolution and MS2 scans collected at 11,250 resolution in profile mode with a maximum injection time of 60 ms, a precursor isolation window of 1.5 m/z with 0.5 m/z offset, and dynamic exclusion set to 5s.

Data processing was performed using MZmine 4.8.30(*36*) and SIRIUS 6.0.7 (*15*) Raw files were converted from Thermo .raw format to .mzML file format and batch processed using the following parameters: mass detection (noise level 1,000), chromatogram building (minimum time span 0.01 min, minimum height 3,000, m/z tolerance 0.1 m/z or 20 ppm), chromatogram deconvolution using the baseline cutoff algorithm (minimum height 31,000, peak duration 0.01–3.0 min, baseline level 300), deisotoping using the isotopic peak grouper (m/z tolerance 0.1 m/z or 20 ppm, RT tolerance 0.1 min, maximum charge 4), peak alignment using the join aligner (m/z tolerance 0.1 m/z or 20 ppm, RT tolerance 0.1 min, m/z weight 75, RT weight 25), and peak filtering (minimum peaks per row 3, minimum peaks per isotope pattern 2).

### Benchmark evaluation on MassSpecGym

#### Inputs

Spectra, reference structures and fold labels were taken from MassSpecGym(*12*). For each MassSpecGym spectrum covered by the HNSW-MS index, the propagation neighborhood was exported as GraphML together with the MS/MS spectra of all neighbors (MGF). In each GraphML, the query was encoded as a fixed seed node carrying only its precursor *m/z*; neighbor nodes carried precursor mass, any library name and structure, and molecular community labels, and edges carried MS/MS cosine similarity and absolute precursor mass difference. Each export was linked to its MassSpecGym record by file-order index, and the link was verified by agreement between the seed precursor mass and the MassSpecGym precursor *m/z*. Fold labels were joined after linking and did not appear in any model-facing file.

#### Evaluation set

The evaluation set comprised randomly selected 18 unique structures with 285 test-fold spectra (of 17,556 in the test fold), corresponding to by first InChIKey block.

#### Blinding and prompt construction

For each query, the name, structure and MassSpecGym accession were removed from all model-facing files; reference structures were stored separately and accessed only at scoring. Neighbors were ranked by an evidence score combining edge cosine similarity, mass residuals consistent with adducts, multimers and water losses, and annotation richness, and the top 25–50 were retained. Each prompt contained the precursor *m/z*, ion mode and peak list of the query; the precursor masses, spectra, structures and query cosine similarities of the neighbors; and a rule-based explanation of neighbor fragments computed offline.

#### Neighbor-only library search

In the library-assisted configuration, the MS/MS spectrum of each neighbor, but never of the query, was reverse-searched against the NIST 2023 MS/MS library with precursor-mass filtering, and matches were supplied to the model as a separate block of evidence. Models were instructed to adopt a library structure only if it was consistent with the precursor mass of the query.

#### Structure assignment

Models received a fixed system prompt, identical for all spectra and not tuned on the benchmark, that required reasoning in the order: mass, network, MS/MS, chemistry and decision, and a single structure was then returned as JSON (SMILES, name, formula, adduct, confidence, rationale and alternatives; detailed prompt is described in Supplementary Materials). Proposed structures had to match the query precursor *m/z* under an even-electron monomer or multimer adduct, including water losses, within 0.05 Da (0.10 Da for multimers). Spectra were processed one at a time, and models were explicitly instructed to not build pack-wide structure indices, run batch mass-fitting scripts or query external registries. Runs that violated this requirement, identifiable by outputs for an entire evaluation set returned within seconds to minutes or by self-reported clustering of spectra by *m/z*, were classed as automated ranking rather than reasoning and discarded.

#### Configurations

Four configurations were evaluated: Grok 4.5 without library search, on all 285 spectra; Claude Opus 4.8 with neighbor-only library search, on 64 spectra, after which the run was stopped because of token cost (approximately 8 million tokens); Qwen3-14B, run locally with neighbor-only library search, on 87 spectra, after which the run reached its runtime limit; and ChemDFM-R without a propagation neighborhood, on all 285 spectra.

#### Scoring

Predictions were scored by first-block InChIKey identity (connectivity), canonical SMILES identity (exact structure), molecular formula identity and Morgan fingerprint Tanimoto similarity (radius 2, 2,048 bits; thresholds 0.7 and 0.85), all computed with RDKit. Where a stored reference InChIKey disagreed with the key computed from the reference SMILES, the computed key was used. Accuracy is reported per spectrum with Wilson 95% confidence intervals, and per unique structure where stated.

### Prediction and confirmation of N-cholyltryptamine

#### Mouse model preparation for the discovery of N-cholyltryptamine

Germ-free (GF) Swiss Webster mice (Taconic) were bred and maintained at Baylor College of Medicine. All procedures were approved by the Baylor College of Medicine Institutional Animal Care and Use Committee and performed in accordance with National Institutes of Health guidelines. Male and female adult mice (3-4 months of age) were randomly assigned to treatments groups and housed in three separate, identical flexible isolators supplied with HEPA-filtered air. All groups were handled identically and maintained on a 12-h light/dark cycle with irradiated food and water available ad libitum. For bifidobacterial colonization, log phase Bifidobacterium species were used. Briefly, Bifidobacterium dentium (ATCC 27678), Bifidobacterium longum subspecies infantis (ATCC 15697), Bifidobacterium breve (ATCC 15698), and Bifidobacterium bifidum (ATCC 29521) were grown anaerobically in Man, Rogosa and Sharpe (MRS) medium (BD, Sparks, MD, USA) at 37°C for 12-14 hrs. The cultures were centrifuged, washed with PBS and combined in equal proportions based on OD₆₀₀ in sterile phosphate-buffered saline (PBS; Gibco). For conventionalization, fecal pellets were obtained from age matched conventional Swiss Webster mice and a fecal slurry was generated in PBS at 100 mg/mL. Mice were gavaged with 200 μL of Bifidobacterium strains or fecal slurry for colonization. Another set of mice were gavaged with sterile PBS to remain germ free. Stool was collected and serial diluted on MRS and BHI-blood agar plates aerobically and anaerobically to examine bacterial colonization. After 3 weeks of colonization, the mice were euthanized and the cecal and fecal samples were collected.

#### Metabolite Extraction and Sample Preparations

For non-targeted metabolomics sample analyses, a 10 µL volume of fecal sample extract was diluted in a 190 µL volume methanol: water (1:1, *v:v*) for a 20-fold dilution, and a 2 µL volume was injected on column for each sample for bioanalysis.

#### Liquid Chromatography – High-Resolution Mass Spectrometry (LC-HRMS) Instrumentation

The LC-HRMS system used for this non-targeted metabolomics study was comprised of a Nexera Series-40 ultrahigh-performance liquid chromatography system outfitted with a SIL-30ACMP autosampler (Shimadzu, Kyoto, Japan) that is coupled to a ZenoTOF 7600+ quadrupole-time-of-flight MS (QTOF MS) system (SCIEX, Framingham, MA, USA). The system was operated using SCIEX OS software (Version 3.3.1.43).

#### Solvents, Supplies, and Consummables

Honeywell – Burdick and Jackson produced LC-MS grade water, methanol, and acetonitrile solvents, and Optima™ LC/MS-grade ammonium formate and formic acid modifiers were purchased from Fisher Scientific (Waltham, MA, USA). The guard and analytical column combination used for reverse-phase, non-targeted metabolomics included an Ultra C18 (10 mm x 2.1 mm; Catalog #: 917450212) guard cartridge coupled to a Raptor C18 (100 mm x 2.1 mm; 2.7-micron; 90Å pore size; Catalog #: 9304A12) purchased from Restek (Bellefonte, PA, USA).

#### Reverse-Phase Positive Mode Liquid Chromatography Conditions

This separation was performed using an Ultra C18 guard and Raptor C18 analytical column combination with mobile phase A (MPA), mobile phase B (MPB) and needlewash (NW) solution preparations consisting of acetonitrile: water (5:95, *v:v*) with 0.1% formic acid, acetonitrile: formic acid (99.9:0.1, *v:v*), and acetonitrile: water (1:1, *v:v*), respectively. After thorough mixing, each solution was degassed using a pulsed bath sonicator (Elmasonic S, Elma Schmidbauer GmbH, Singen, Germany) for 15 minutes (min) without heating prior to use. These solutions were stored at ambient temperature and expired two months after preparation.

The LC system program was specified to deliver a mobile phase flowrate of 0.300 mL/min at an initial proportion of MPA: MPB (100:0, *v:v*), the analytical column was heated to 50°C, and the mobile phase gradient program was specified as follows: 0.0-2.0 min, 0% MPB; 2.0-11.0 min, 0-95% MPB; 11.0-13.0 min, 95% MPB; 13.0-13.01 min, 95-0% MPB; and, 13.01-15.0 min, 0% MPB – with a per sample cycle time of ∼15.5 min. The SIL-30ACMP autosampler settings were specified as follows: Sampling speed, 5.0 µL/second (s); Cooler temperature, +15°C; Temperature control, enabled for Rack plate L, Rack plate M, and Rack plate R; Use air gap volume, No; Rinse type, External only; Rinse mode, Before and after aspiration; Dip time: 1 s; Rinse method, Rinse port only; Time, 2 s; Sampling speed, 35 µL/s; Rinse volume, 500 µL; Measuring line purge volume, 100 µL.

#### Non-targeted metabolomics LC-MS/MS method

The ZenoTOF 7600+ was outfitted with an OptiFlow Pro Ionization Source with a 50-200 µL/min electrospray ionization (ESI) probe installed. Global mass spec parameters were specified as follows: Workflow, Small molecule; Intact protein mode, False; Method duration, 15 min; Total scan time, 0.566 s (software determined); Estimated cycles, 1,589 (software determined). The Source and Gas Parameters were specified as: Ion source gas 1, 35 psi; Ion source gas 2, 70 psi; Curtain gas, 40 psi; collision-activated dissociation (CAD) gas, 9; Temperature, 450°C.

The experimental settings were specified as: Scan setting, Information-Dependent Acquisition (IDA); Polarity, Positive; Spray voltage, +5,500 volts (V); TOF start mass, 70 Dalton (Da); TOF stop mass, 1500 Da; Accumulation time, 0.1 seconds (s); Declustering potential (DP), +80 V; DP spread, +/-0 V; Collision Energy (CE), +12 electron-volts (eV); CE spread, +/-0 eV. Advanced Experimental Settings were specified as: Time bins to sum, 4; Channels 1-4, enabled; Manual QJet radio-frequency (RF) amplitude, disabled; QJet RF amplitude, 193.788030257723 V (software determined). IDA Criteria were specified as: Setting, Small molecule; Maximum candidate ions, 40; Intensity threshold exceeds (counts/s), 250 cps; Dynamic background subtraction, enabled; Exclude former candidate ions, True (enabled for 6 sec after 3 occurrences); Dynamic CE for MS/MS, False; Dynamic ETC for MS/MS, False; Charge state, False; and, Isotope to select, Monoisotopic. Advanced IDA criteria were specified as: Exclude Isotope, False; Mass tolerance (+/-mDa), 50; Candidate mass range, False; Adjust CE when using iTRAQ reagent, False; Inclusion list, False; Exclusion list, False; Mass Defect Filter, False; Isotope Matching, False; and, Neutral Loss, False.

The TOF MSMS settings were specified as: Polarity, Positive; Spray Voltage, +5500 V; Fragmentation mode, collisionally-induced dissociation (CID); TOF start mass, 50 Da; TOF stop mass, 1500 Da; Accumulation time, 0.005 s; DP, +80 V; DP spread, +/-0 V; CE, +50 eV; and, CE spread, +/-10 eV. Advanced Experimental Settings were specified as: Q1 resolution, Unit; Zeno pulsing, True; Zeno threshold (CPS), 20,000; Time bins to sum, 4; Channels 1-4, enabled; Manual QJet RF amplitude, disabled; QJet RF amplitude, 202.72 V.

#### Query spectrum and blinding

The query was a single positive-mode spectrum. The query spectrum was not searched against the NIST 2023 MS/MS library, while the names and NIST matches of the neighbors were retained, as in the MassSpecGym evaluation. The precursor was measured at m/z 551.382, which lies 2.33 mDa below the theoretical [M+H]⁺ of 551.38433 for C₃₄H₅₀N₂O₄.

#### Propagation neighborhood

The neighborhood was drawn from a 459-node subnetwork and comprised the query node, its 25 first-order neighbors and 40 annotated two-hop nodes, the latter supplied as name and precursor mass only, without MS/MS, NIST matches or formula assignments. Edge cosines across the ego layer ranged from 0.763 to 0.774.

The neighborhood was submitted independently to Grok 4.6, Opus 5.5 and Grok 4.7 under the standard prompt (Methods, Reasoning models and inference protocol). All three returned the same connectivity (InChIKey first block SCTPNCMIQYWHOJ) at confidences of 0.76, 0.60 and 0.77; Grok 4.6 and Opus 5.5 returned the full InChIKey SCTPNCMIQYWHOJ-LXCBOGQWSA-N, and Grok 4.7 left stereochemistry unassigned. Alternatives retained at low confidence comprised a 3,6,7-trihydroxy isomer (0.15 by Opus 5.5, 0.34 by Grok 4.7), a 7β isomer (0.05 by Opus 5.5) and serotonin-deoxycholic acid (0.03 by Opus 5.5). For comparison, SIRIUS 5.8.6 ranked C₂₉H₅₀N₄O₆ first for the same spectrum, with a score of 43.8, an explained intensity of 0.89 and 15 explained fragments, at a theoretical m/z of 551.38031; all three models set this formula aside. Both Grok calls described a one water loss ion that the query spectrum does not contain, and the ions at m/z 89.058, 133, 745.405 and 789.443 were treated by Opus 5.5 as co-isolated background.

#### Synthesis of N-cholyltryptamine

The standard was synthesized to confirm the structure of a feature suspected to be N-cholyltryptamine based on its MS/MS fragmentation reproducibility. Deoxycholic acid (0.125 mmol, 1.0 equiv) was dissolved in anhydrous tetrahydrofuran (2.5 mL) in a 20 mL vial at 0 °C. Ethyl chloroformate (0.15 mmol, 1.2 equiv) and triethylamine (0.15 mmol, 1.2 equiv) were added in that order, and the mixture was stirred at 0 °C for 2.5 h. Tryptamine (0.193 mmol, 1.54 equiv) in aqueous sodium bicarbonate (0.188 mmol, 1.50 equiv; 2.5 mL) was then added. Stirring was continued at 0 °C for 4 h and then for 2 h while the mixture warmed to room temperature. The crude product was dried under vacuum, washed with water, and stored at −80 °C until use.

#### Mass spectrometry of the standard and the query

The synthetic standard was acquired on an Orbitrap Exploris 240 operated in FTMS mode at HCD 40 as described above for the test library data collection, with the precursor observed at m/z 551.3843, 0.03 mDa from theory, the MS1 peak at 6.10 min and the MS2 scan at 6.11 min.

### Reasoning models and inference protocol

#### Models

Propagation neighborhoods were interpreted by two frontier reasoning models accessed through their providers’ APIs, Claude Opus 4.8 (Anthropic) and Grok 4.5 (xAI), between August and October 2026. On MassSpecGym, two further models were evaluated for comparison: Qwen3-14B, an open-weight model run locally (via Ollama), and ChemDFM-R, a chemistry reasoning model given the query spectrum without a propagation neighborhood. Sampling settings were: temperature, maximum output tokens, reasoning budget. No model was trained, fine-tuned or otherwise adapted to the task.

#### Prompt

Each query was submitted as a single prompt consisting of a fixed system instruction and a query-specific evidence block. Specific prompts instruction, identical for all queries and models (verbatim in Supplementary Materials), defined the task as network-assisted structure annotation and set five rules. (i) Use only peaks written in the prompt. (ii) Mass first: a candidate structure must reproduce the observed precursor *m/z* as an even-electron ion, either a monomer or a multimer adduct, allowing single or multiple water losses, within 0.05 Da (0.10 Da for multimers). (iii) prefer higher order neighbors whose annotated structures relate to the query by plausible mass differences are the primary source of structural hypotheses. (iv) Fragmentation evidence, including query fragments and the spectral similarity of the query to each neighbor (within 0.5 Da) is used to discriminate among candidates. (v) Reasoning proceeds in a fixed order: mass, network, MS/MS, chemistry and decision. The model returned a single JSON object containing the proposed structure (SMILES), name, molecular formula, adduct, a confidence score, a brief rationale and ranked alternatives.

#### Execution

Each query was processed in a separate request, with no access to other queries, external databases or the internet. Models could not write or run code that operated across the evaluation set. Each query received a single pass, without retries, prompt revision or selection among outputs. Returned SMILES were parsed with RDKit, and responses without a valid structure were scored as incorrect. Wall-clock time and token use were recorded for every run, and runs inconsistent with per-query reasoning were excluded (see Benchmark evaluation on MassSpecGym). The confidence score was taken as reported by the model on a 0-100% scale. The evidence message is assembled using eleven separate parameters. (i) Query header: precursor m/z, the hidden query-node identifier, degree, the count of neighbors within 0.5 Da, the count of multimer-consistent neighbors, whether MS/MS is available, and retention time. (ii) Query MS/MS: the quality line, the 15 most intense peaks as m/z and percent of the base peak, retention time, and ion mode. (iii) An offline MS/MS explanation listing neutral losses from the precursor and compares the query with the three neighbors of highest MS/MS cosine (explanation independent of SIRIUS). (iv) Spectral library hits for the query. (v) Up to six approximate monomer masses for the case in which the precursor is a multimer. (vi) Near-isobar neighbors, multimer-consistent neighbors, and mass-consistent library structures. (vii) The 25 direct neighbors in the combined cosine order. (viii) The selected second-hop nodes. (ix) The SIRIUS formula block, when that experiment includes formula hypotheses (optional). (x) The instruction to fit the mass, to use the fragments to choose among mass-consistent candidates, and to return the JSON object. (xi) Neighbor library hits, when that experiment uses library search.

#### Comparison between models

The frontier models were run independently on identical prompts. If/when used for prediction of the same query, they were considered to agree when their predictions shared the first layer of InChIKey with stereochemistry removed. Accuracy was computed separately for queries on which the models agreed and on which they disagreed. The union of correct predictions across both models is reported as an upper bound on what a consensus scheme could recover.

## Acknowledgments

A.A.A. was supported by Agilent. The Texas Children’s Research Institute (TCRI) provides salary support to the staff of the Virginia and L.E. Simmons Family Foundation Mass Spectrometry Laboratory housed within the TCRI-Microbiome Center, and purchased all reagents, consumables and durable supplies.

## Contributions

A.A.A. formulated the concept of molecular neighborhood annotation propagation. A.S. and V.B. devised the mathematical underpinning. A.S. performed code engineering for the generation of propagation neighborhood networks and generated propagation neighborhood networks from experimental data. A.A.A., A.M., D.M., A.R. curated the data used in the analyses. B.O. carried out global network connectivity statistics. T.D.H., M.A.E. collected data for the humanized mouse model and gut bioreactor. N.S., S.D., M.P. conducted organic synthesis for novel structures. A.M. performed reference data collection. A.M. and P.H. conducted AI-based annotation propagation/unknown prediction. A.A.A. provided supervision for the project. A.A.A., A.R., E.A.C., V.B., and A.S. wrote and edited the manuscript. All authors reviewed and approved the manuscript.

## Data/code access

We have deposited the code at GitHub: github.com/Melin2012/ego-mol-llm, Apache-2.0. The code is open source and released under an LGPL-3 license. Any additional information required to reanalyze the data reported in this study is available from the corresponding author upon request.

## Supplementary Materials

https://docs.google.com/spreadsheets/d/1BOBhd8e4LehX4Sowsw3NRH9nwj4emrewASbG0LyMYiM/edit?usp=sharing

**Supplemental Table 1.** Prediction accuracy on 2513 spectra from Arome’s 900 standards

### Prompt

Each query is one request that contains a system instruction, identical for every spectrum, and one evidence message. The system instruction tells the model to assign a neutral structure from the neighborhood, the fragmentation, and any optional library or formula block. The unknown may be absent from the evidence. The rules, in order, are as follows.

1. Use only peaks that are given in the prompt.
2. Mass first. The structure must fit the precursor as an even-electron monomer or multimer adduct, including water loss. The listed adducts include [M+H]+, [M+Na]+, [M+K]+, [M+NH4]+, [M+H-H2O]+, the corresponding negative ions, and the dimer and trimer forms [2M+H]+, [2M+Na]+, [2M-H]-, and [3M+H]+.
3. Prefer a neighbor that is high on both the network cosine and the MS/MS cosine. A neighbor or a library hit is copied only when that structure fits the precursor mass and the query fragments. A new structure that fits both is allowed.
4. A neighbor within 0.5 Da is evidence for a monomer, and is set aside when its formula cannot fit the precursor. A dimer or trimer is used when the structure mass fits.
5. Reason in the fixed order mass, network, MS/MS, chemistry, decision. End with one JSON object and nothing after it.

The JSON object contains the neutral SMILES, a name, the molecular formula, the adduct of the observed ion, a confidence from 0 to 1, a rationale of five to ten sentences, and alternatives. The SMILES must be valid in RDKit.

The evidence message is assembled in the following order.

1. Query header: precursor m/z, the hidden query-node identifier, degree, the count of neighbors within 0.5 Da, the count of multimer-consistent neighbors, whether MS/MS is available, and retention time.
2. Query MS/MS: the quality line, the 15 most intense peaks as m/z and percent of the base peak, retention time, and ion mode.
3. An offline MS/MS explanation. It lists neutral losses from the precursor and compares the query with the three neighbors of highest MS/MS cosine. This explanation is independent of SIRIUS.
4. Spectral library hits for the query. This block is empty. The query spectrum is not searched.
5. Up to six approximate monomer masses for the case in which the precursor is a multimer.
6. Near-isobar neighbors, multimer-consistent neighbors, and mass-consistent library structures.
7. The 25 direct neighbors in the combined cosine order.
8. The selected second-hop nodes.
9. The SIRIUS formula block, when that experiment includes formula hypotheses. Otherwise this block is omitted.
10. The instruction to fit the mass, to use the fragments to choose among mass-consistent candidates, and to return the JSON object.
11. Neighbor library hits, when that experiment uses library search.

Comparisons that isolate one source of evidence use the same prompt with the corresponding block removed. Network-only reasoning omits the query peak list.

Spectrum-only reasoning omits the neighborhood. Formula-assisted reasoning inserts the SIRIUS block described below. A synthetic-compound experiment can add an instruction that biological frequency is not a prior. No experiment in this study tunes the system instruction on the spectra being scored.

## References and Notes

1. D. J. Newman, G. M. Cragg, Natural Products as Sources of New Drugs over the Nearly Four Decades from 01/1981 to 09/2019. J Nat Prod 83, 770–803 (2020).

2. J. Jumper, R. Evans, A. Pritzel, T. Green, M. Figurnov, O. Ronneberger, K. Tunyasuvunakool, R. Bates, A. Žídek, A. Potapenko, A. Bridgland, C. Meyer, S. A. A. Kohl, A. J. Ballard, A. Cowie, B. Romera-Paredes, S. Nikolov, R. Jain, J. Adler, T. Back, S. Petersen, D. Reiman, E. Clancy, M. Zielinski, M. Steinegger, M. Pacholska, T. Berghammer, S. Bodenstein, D. Silver, O. Vinyals, A. W. Senior, K. Kavukcuoglu, P. Kohli, D. Hassabis, Highly accurate protein structure prediction with AlphaFold. Nature 596, 583–589 (2021).

3. A. A. Aksenov, I. Laponogov, Z. Zhang, S. L. F. Doran, I. Belluomo, D. Veselkov, W. Bittremieux, L. F. Nothias, M. Nothias-Esposito, K. N. Maloney, B. B. Misra, A. V. Melnik, A. Smirnov, X. Du, K. L. Jones 2nd, K. Dorrestein, M. Panitchpakdi, M. Ernst, J. J. J. van der Hooft, M. Gonzalez, C. Carazzone, A. Amézquita, C. Callewaert, J. T. Morton, R. A. Quinn, A. Bouslimani, A. A. Orio, D. Petras, A. M. Smania, S. P. Couvillion, M. C. Burnet, C. D. Nicora, E. Zink, T. O. Metz, V. Artaev, E. Humston-Fulmer, R. Gregor, M. M. Meijler, I. Mizrahi, S. Eyal, B. Anderson, R. Dutton, R. Lugan, P. L. Boulch, Y. Guitton, S. Prevost, A. Poirier, G. Dervilly, B. Le Bizec, A. Fait, N. S. Persi, C. Song, K. Gashu, R. Coras, M. Guma, J. Manasson, J. U. Scher, D. K. Barupal, S. Alseekh, A. R. Fernie, R. Mirnezami, V. Vasiliou, R. Schmid, R. S. Borisov, L. N. Kulikova, R. Knight, M. Wang, G. B. Hanna, P. C. Dorrestein, K. Veselkov, Auto-deconvolution and molecular networking of gas chromatography-mass spectrometry data. Nat Biotechnol 39, 169–173 (2021).

4. R. R. da Silva, P. C. Dorrestein, R. A. Quinn, Illuminating the dark matter in metabolomics. Proc. Natl. Acad. Sci. U. S. A. 112, 12549–12550 (2015).

5. M. Wang, J. J. Carver, V. V. Phelan, L. M. Sanchez, N. Garg, Y. Peng, D. D. Nguyen, J. Watrous, C. A. Kapono, T. Luzzatto-Knaan, C. Porto, A. Bouslimani, A. V. Melnik, M. J. Meehan, W.-T. Liu, M. Crüsemann, P. D. Boudreau, E. Esquenazi, M. Sandoval-Calderón, R. D. Kersten, L. A. Pace, R. A. Quinn, K. R. Duncan, C.-C. Hsu, D. J. Floros, R. G. Gavilan, K. Kleigrewe, T. Northen, R. J. Dutton, D. Parrot, E. E. Carlson, B. Aigle, C. F. Michelsen, L. Jelsbak, C. Sohlenkamp, P. Pevzner, A. Edlund, J. McLean, J. Piel, B. T. Murphy, L. Gerwick, C.-C. Liaw, Y.-L. Yang, H.-U. Humpf, M. Maansson, R. A. Keyzers, A. C. Sims, A. R. Johnson, A. M. Sidebottom, B. E. Sedio, A. Klitgaard, C. B. Larson, C. A. B. P D. Torres-Mendoza, D. J. Gonzalez, D. B. Silva, L. M. Marques, D. P. Demarque, E. Pociute, E. C. O’Neill, E. Briand, E. J. N. Helfrich, E. A. Granatosky, E. Glukhov, F. Ryffel, H. Houson, H. Mohimani, J. J. Kharbush, Y. Zeng, J. A. Vorholt, K. L. Kurita, P. Charusanti, K. L. McPhail, K. F. Nielsen, L. Vuong, M. Elfeki, M. F. Traxler, N. Engene, N. Koyama, O. B. Vining, R. Baric, R. R. Silva, S. J. Mascuch, S. Tomasi, S. Jenkins, V. Macherla, T. Hoffman, V. Agarwal, P. G. Williams, J. Dai, R. Neupane, J. Gurr, A. M. C. Rodríguez, A. Lamsa, C. Zhang, K. Dorrestein, B. M. Duggan, J. Almaliti, P.-M. Allard, P. Phapale, L.-F. Nothias, T. Alexandrov, M. Litaudon, J.-L. Wolfender, J. E. Kyle, T. O. Metz, T. Peryea, D.-T. Nguyen, D. VanLeer, P. Shinn, A. Jadhav, R. Müller, K. M. Waters, W. Shi, X. Liu, L. Zhang, R. Knight, P. R. Jensen, B. O. Palsson, K. Pogliano, R. G. Linington, M. Gutiérrez, N. P. Lopes, W. H. Gerwick, B. S. Moore, P. C. Dorrestein, N. Bandeira, Sharing and community curation of mass spectrometry data with Global Natural Products Social Molecular Networking. Nat. Biotechnol. 34, 828–837 (2016).

6. M. Wang, A. K. Jarmusch, F. Vargas, A. A. Aksenov, J. M. Gauglitz, K. Weldon, D. Petras, R. da Silva, R. Quinn, A. V. Melnik, J. J. J. van der Hooft, A. M. Caraballo-Rodríguez, L. F. Nothias, C. M. Aceves, M. Panitchpakdi, E. Brown, F. Di Ottavio, N. Sikora, E. O. Elijah, L. Labarta-Bajo, E. C. Gentry, S. Shalapour, K. E. Kyle, S. P. Puckett, J. D. Watrous, C. S. Carpenter, A. Bouslimani, M. Ernst, A. D. Swafford, E. I. Zúñiga, M. J. Balunas, J. L. Klassen, R. Loomba, R. Knight, N. Bandeira, P. C. Dorrestein, Mass spectrometry searches using MASST. Nat Biotechnol 38, 23–26 (2020).

7. M. Mongia, T. M. Yasaka, Y. Liu, M. Guler, L. Lu, A. Bhagwat, B. Behsaz, M. Wang, P. C. Dorrestein, H. Mohimani, Fast mass spectrometry search and clustering of untargeted metabolomics data. Nat Biotechnol 42, 1672–1677 (2024).

8. H. Zhang, Q. Yang, T. Xie, Y. Wang, Z. Zhang, H. Lu, MSBERT: Embedding Tandem Mass Spectra into Chemically Rational Space by Mask Learning and Contrastive Learning. Anal Chem 96, 16599–16608 (2024).

9. R. Bushuiev, A. Bushuiev, R. Samusevich, C. Brungs, J. Sivic, T. Pluskal, Self-supervised learning of molecular representations from millions of tandem mass spectra using DreaMS. Nat Biotechnol 44, 630–640 (2026).

10. S. Goldman, J. Wohlwend, M. Stražar, G. Haroush, R. J. Xavier, C. W. Coley, Annotating metabolite mass spectra with domain-inspired chemical formula transformers. *Nat*. Mach. Intell. 5, 965–979 (2023).

11. Z. A. Quinlan, I. Koester, A. T. Aron, D. Petras, L. I. Aluwihare, P. C. Dorrestein, C. E. Nelson, L. Wegley Kelly, ConCISE: Consensus Annotation Propagation of Ion Features in Untargeted Tandem Mass Spectrometry Combining Molecular Networking and Metabolite Structure Prediction. Metabolites 12 (2022).

12. R. Bushuiev, A. Bushuiev, N. F. de Jonge, A. Young, F. Kretschmer, R. Samusevich, J. Heirman, F. Wang, L. Zhang, K. Dührkop, M. Ludwig, N. A. Haupt, A. Kalia, C. Brungs, R. Schmid, R. Greiner, B. Wang, D. S. Wishart, L.-P. Liu, J. Rousu, W. Bittremieux, H. Rost, T. D. Mak, S. Hassoun, F. Huber, J. J. J. van der Hooft, M. A. Stravs, S. Böcker, J. Sivic, T. Pluskal, MassSpecGym: A benchmark for the discovery and identification of molecules (2024). http://arxiv.org/abs/2410.23326.

13. H. Liu, R. Bushuiev, I. Lightheart, M. Manjrekar, A. Bushuiev, M. Lederbauer, F. Jozefov, Y. Wang, S. Hassoun, J. Sivic, J. Taylor, R. Wang, D. Healey, T. Pluskal, C. W. Coley, MassSpecGym in the Wild: Uncovering and Correcting Evaluation Pitfalls in AI-Driven Molecule Discovery (2026). http://arxiv.org/abs/2606.19624.

14. K. Dührkop, H. Shen, M. Meusel, J. Rousu, S. Böcker, Searching molecular structure databases with tandem mass spectra using CSI:FingerID. Proceedings of the National Academy of Sciences 112, 12580–12585 (2015).

15. K. Dührkop, M. Fleischauer, M. Ludwig, A. A. Aksenov, A. V. Melnik, M. Meusel, P. C. Dorrestein, J. Rousu, S. Böcker, SIRIUS 4: a rapid tool for turning tandem mass spectra into metabolite structure information. Nature Methods 16, 299–302 (2019).

16. H. Wang, H. Zhang, Z.-J. Zhu, Structure-informed deep generation enables de novo metabolite annotation in untargeted metabolomics. Nature Communications 17, 5426 (2026).

17. E. A. Coler, A. Melnik, A. Lotfi, D. Moradi, B. Ahiadu, P. W. Portal Gomes, A. Patan, V. Charron-Lamoureux, P. C. Dorrestein, S. Barnes, V. Boginski, A. Semenov, A. A. Aksenov, Ordering molecular diversity in untargeted metabolomics via molecular community networking. Cell Rep Methods 6, 101468 (2026).

18. A. Semenov, A. M. P. Roberts, E. A. Coler, S. Gupta, E. Kopylova, A. V. Melnik, V. Boginski, A. A. Aksenov, Real-time repository-scale spectral search and global molecular networking with HNSW-MS, bioRxiv (2026). 10.64898/2026.06.02.729602.

19. W. Bittremieux, N. E. Avalon, S. P. Thomas, S. A. Kakhkhorov, A. A. Aksenov, P. W. P. Gomes, C. M. Aceves, A. M. Caraballo-Rodríguez, J. M. Gauglitz, W. H. Gerwick, T. Huan, A. K. Jarmusch, R. F. Kaddurah-Daouk, K. B. Kang, H. W. Kim, T. Kondić, H. Mannochio-Russo, M. J. Meehan, A. V. Melnik, L.-F. Nothias, C. O’Donovan, M. Panitchpakdi, D. Petras, R. Schmid, E. L. Schymanski, J. J. J. van der Hooft, K. C. Weldon, H. Yang, S. Xing, J. Zemlin, M. Wang, P. C. Dorrestein, Open access repository-scale propagated nearest neighbor suspect spectral library for untargeted metabolomics. Nat Commun 14, 8488 (2023).

20. X. Wang, A. Patan, H. N. Zhao, V. Charron-Lamoureux, Y. Shin, D. Petras, Y. Hong, B. P. Bowen, T. R. Northen, P. C. Dorrestein, M. Wang, Agentic AI for Structural Elucidation and Discovery of Drug Metabolites from Mass Spectrometry Data, bioRxiv (2026). 10.64898/2026.06.23.734138.

21. E. A. Coler, W. Chen, A. V. Melnik, J. T. Morton, A. A. Aksenov, Metabolomics in the era of artificial intelligence. Microbiota Host 2 (2024).

22. R. A. Quinn, A. V. Melnik, A. Vrbanac, T. Fu, K. A. Patras, M. P. Christy, Z. Bodai, P. Belda-Ferre, A. Tripathi, L. K. Chung, M. Downes, R. D. Welch, M. Quinn, G. Humphrey, M. Panitchpakdi, K. C. Weldon, A. Aksenov, R. da Silva, J. Avila-Pacheco, C. Clish, S. Bae, H. Mallick, E. A. Franzosa, J. Lloyd-Price, R. Bussell, T. Thron, A. T. Nelson, M. Wang, E. Leszczynski, F. Vargas, J. M. Gauglitz, M. J. Meehan, E. Gentry, T. D. Arthur, A. C. Komor, O. Poulsen, B. S. Boland, J. T. Chang, W. J. Sandborn, M. Lim, N. Garg, J. C. Lumeng, R. J. Xavier, B. I. Kazmierczak, R. Jain, M. Egan, K. E. Rhee, D. Ferguson, M. Raffatellu, H. Vlamakis, G. G. Haddad, D. Siegel, C. Huttenhower, S. K. Mazmanian, R. M. Evans, V. Nizet, R. Knight, P. C. Dorrestein, Global chemical effects of the microbiome include new bile-acid conjugations. Nature 579, 123–129 (2020).

23. O. Fiehn, D. Robertson, J. Griffin, M. van der Werf, B. Nikolau, N. Morrison, L. W. Sumner, R. Goodacre, N. W. Hardy, C. Taylor, J. Fostel, B. Kristal, R. Kaddurah-Daouk, P. Mendes, B. van Ommen, J. C. Lindon, S.-A. Sansone, The metabolomics standards initiative (MSI). Metabolomics 3, 175–178 (2007).

24. B. Rimal, S. L. Collins, C. E. Tanes, E. R. Rocha, M. A. Granda, S. Solanki, N. J. Hoque, E. C. Gentry, I. Koo, E. R. Reilly, F. Hao, D. Paudel, V. Singh, T. Yan, M. S. Kim, K. Bittinger, J. P. Zackular, K. W. Krausz, D. Desai, S. Amin, J. P. Coleman, Y. M. Shah, J. E. Bisanz, F. J. Gonzalez, J. P. Vanden Heuvel, G. D. Wu, B. S. Zemel, P. C. Dorrestein, E. E. Weinert, A. D. Patterson, Bile salt hydrolase catalyses formation of amine-conjugated bile acids. Nature 626, 859–863 (2024).

25. M. W. Mullowney, A. Fiebig, M. K. Schnizlein, M. McMillin, A. R. Rose, J. Koval, D. Rubin, S. Dalal, M. L. Sogin, E. B. Chang, A. M. Sidebottom, S. Crosson, Microbially catalyzed conjugation of GABA and tyramine to bile acids. Journal of Bacteriology, doi: 10.1128/jb.00426-23 (2024).

26. B. B. Williams, A. H. Van Benschoten, P. Cimermancic, M. S. Donia, M. Zimmermann, M. Taketani, A. Ishihara, P. C. Kashyap, J. S. Fraser, M. A. Fischbach, Discovery and characterization of gut microbiota decarboxylases that can produce the neurotransmitter tryptamine. Cell Host Microbe 16, 495–503 (2014).

27. Y. Bhattarai, B. B. Williams, E. J. Battaglioli, W. R. Whitaker, L. Till, M. Grover, D. R. Linden, Y. Akiba, K. K. Kandimalla, N. C. Zachos, J. D. Kaunitz, J. L. Sonnenburg, M. A. Fischbach, G. Farrugia, P. C. Kashyap, Gut Microbiota-Produced Tryptamine Activates an Epithelial G-Protein-Coupled Receptor to Increase Colonic Secretion. Cell Host Microbe 23, 775–785.e5 (2018).

28. A. Wahlström, S. I. Sayin, H.-U. Marschall, F. Bäckhed, Intestinal Crosstalk between Bile Acids and Microbiota and Its Impact on Host Metabolism. Cell Metab 24, 41–50 (2016).

29. S. L. Collins, J. G. Stine, J. E. Bisanz, C. D. Okafor, A. D. Patterson, Bile acids and the gut microbiota: metabolic interactions and impacts on disease. Nat Rev Microbiol 21, 236–247 (2023).

30. A. Agus, J. Planchais, H. Sokol, Gut Microbiota Regulation of Tryptophan Metabolism in Health and Disease. Cell Host Microbe 23, 716–724 (2018).

31. H. M. Roager, T. R. Licht, Microbial tryptophan catabolites in health and disease. Nat Commun 9, 3294 (2018).

32. D. L. Murphy, L. Tamarkin, N. A. Garrick, P. L. Taylor, S. P. Markey, “Trace indoleamines in the central nervous system” in Neuropsychopharmacology of the Trace Amines (Humana Press, Totowa, NJ, 1985), pp. 343–360.

33. D. V. Guzior, M. Okros, M. Shivel, B. Armwald, C. Bridges, Y. Fu, C. Martin, A. L. Schilmiller, W. M. Miller, K. M. Ziegler, M. D. Sims, M. E. Maddens, S. F. Graham, R. P. Hausinger, R. A. Quinn, Bile salt hydrolase acyltransferase activity expands bile acid diversity. Nature 626, 852–858 (2024).

34. N. G. Mahieu, G. J. Patti, Systems-Level Annotation of a Metabolomics Data Set Reduces 25 000 Features to Fewer than 1000 Unique Metabolites. Anal Chem 89, 10397–10406 (2017).

35. M. Bohde, H. Liu, M. Manjrekar, M. Lederbauer, S. Ji, R. Wang, C. W. Coley, FRIGID: Scaling Diffusion-Based Molecular Generation from Mass Spectra at Training and Inference Time (2026). http://arxiv.org/abs/2604.16648.

36. R. Schmid, S. Heuckeroth, A. Korf, A. Smirnov, O. Myers, T. S. Dyrlund, R. Bushuiev, K. J. Murray, N. Hoffmann, M. Lu, A. Sarvepalli, Z. Zhang, M. Fleischauer, K. Dührkop, M. Wesner, S. J. Hoogstra, E. Rudt, O. Mokshyna, C. Brungs, K. Ponomarov, L. Mutabdžija, T. Damiani, C. J. Pudney, M. Earll, P. O. Helmer, T. R. Fallon, T. Schulze, A. Rivas-Ubach, A. Bilbao, H. Richter, L.-F. Nothias, M. Wang, M. Orešič, J.-K. Weng, S. Böcker, A. Jeibmann, H. Hayen, U. Karst, P. C. Dorrestein, D. Petras, X. Du, T. Pluskal, Integrative analysis of multimodal mass spectrometry data in MZmine 3. Nature Biotechnology 41, 447–449 (2023).

37. D. Schnable, N. Chaudhri, R. Li, M. Zeller, C. Brückner, Evaluation of Octaethyl-7,17-dioxobacteriochlorin as a Ligand for Transition Metals. Inorg Chem 59, 2870–2880 (2020).

38. S. Vaidya, M. Sharma, C. Brückner, R. M. Kasi, Rhodamine-Installed Polynorbornenes: Molecular Design, Structure, and Stimuli-Responsive Properties. ACS Omega 6, 15017–15028 (2021).

39. C. Brückner, N. Chaudhri, D. E. Nevonen, S. Bhattacharya, A. Graf, E. Kaesmann, R. Li, M. J. Guberman-Pfeffer, T. Mani, A. Nimthong-Roldán, M. Zeller, A. A. P. Chauvet, V. Nemykin, Structural and Photophysical Characterization of All Five Constitutional Isomers of the Octaethyl-β,β’-dioxo-bacterio- and -isobacteriochlorin Series. Chemistry 27, 16189–16203 (2021).

40. N. Chaudhri, M. J. Guberman-Pfeffer, R. Li, M. Zeller, C. Brückner, β-Trioxopyrrocorphins: pyrrocorphins of graded aromaticity. Chem Sci 12, 12292–12301 (2021).

41. Y. Blériot, C. R. Veighey, K. H. Smelt, J. Cadefau, W. Stalmans, K. Biggadike, A. L. Lane, M. Müller, D. J. Watkin, G. W. J. Fleet, The first example of a: Inhibition of glucokinase. Tetrahedron Asymmetry 7, 2761–2772 (1996).

42. Y. Blériot, C. F. Masaguer, J. Charlwood, B. G. Winchester, A. L. Lane, S. Crook, D. J. Watkin, G. W. J. Fleet, 6R-, and 6S, −6C-methylglucose from D-glucuronolactone: Efficient synthesis of a seven carbon fucose analogue: Inhibition of some enzymes of primary metabolism. Tetrahedron 53, 15135–15146 (1997).

43. R. U. Lemieux, T. C. Wong, H. Thøgersen, The synthesis and conformational properties of the diastereoisomeric βDGal(1→4)βDGlcNAc(1→6)6-C-CH3-D-Gal trisaccharides. Can. J. Chem. 60, 81–86 (1982).

44. C. Lough, O. Hindsgaul, R. U. Lemieux, Synthesis and conformational properties of methyl 6,6-di-C-methyl-beta-D-galactopyranoside. Probes for the combining sites of D-galactosyl-binding proteins. Carbohydr Res 120, 43–53 (1983).

45. A. Y. Plodukhin, K. Hernández, S. Demirovic, T. Ogunmilade, B. Q. Mercado, M. Kennedy, A. Angeles-Boza, A. Planas, M. W. Peczuh, Unnatural C6-Substituted d-Galactosides as Attenuated Substrates of Glycosyl Hydrolases. Chembiochem 27, e70524 (2026).

